# Piggybacking on the Cholera Toxin: Identification of a Toxoid-binding Protein as an Approach for Targeted Delivery of Proteins to Motor Neurons

**DOI:** 10.1101/2020.05.11.982132

**Authors:** Matthew R. Balmforth, Jessica Haigh, Christian Tiede, Darren C. Tomlinson, Jim Deuchars, Michael E. Webb, W. Bruce Turnbull

## Abstract

A significant unmet need exists for the delivery of biologic drugs such as polypeptides or nucleic acids, to the central nervous system (CNS) for the treatment and understanding of neurodegenerative diseases. Naturally occurring toxoids have been considered as tools to meet this need. However, due to the complexity of tethering macromolecular drugs to toxins, and the inherent dangers of working with large quantities of recombinant toxin, no such route has been successfully exploited. Developing a method where toxoid and drug can be assembled immediately prior to *in vivo* administration has the potential to circumvent some of these issues. Using a phage-display screen, we identified two antibody mimetics, Anti-Cholera Toxoid Affimer (ACTA) -A2 and ACTA-C6 that non-covalently associate with the non-binding face of the cholera toxin B-subunit. In a first step toward the development of a non-viral motor neuron drug-delivery vehicle, we show that Affimers can be selectively delivered to motor neurons *in vivo*.

## INTRODUCTION

Research into treatments for degenerative diseases of motor neurons such as amyotrophic lateral sclerosis (ALS), is hampered in part due to the considerable challenges associated with delivery of molecular probes to the central nervous system (CNS).^1^ To complicate matters, recent trends towards the use of biologics as drugs have resulted in a shift in focus from small molecules to macromolecules such as nucleic acids and antibodies.^2^ Typical strategies for cellular penetration and macromolecule delivery have relied in the past on the use of virally derived cell-penetrating peptides (CPPs).^3–5^ But whilst efficient, these vectors lack specificity in both cell targeting and mechanism of cell entry.^6^ Viral delivery methods, such as those based on the Herpes Simplex virus, have been used successfully to transfer therapeutic genes into the CNS for treatment of conditions such as ALS and spinal cord injury.^7–9^ Viral-mediated gene therapy has been limited however by concerns over vector safety and expense of production.^10^ A significant need therefore exists for the development of novel technologies to deliver biologics into motor neurons via peripheral administration routes.

AB_5_ proteins represent a class of bacterial toxins that gain entry to certain cell types through interactions with highly specific surface markers to trigger receptor-mediated endocytosis.^11–13^ One toxoid from the AB_5_ family, cholera toxin B-subunit (CTB), binds to ganglioside GM1, a membrane-tethered carbohydrate highly expressed at the neuromuscular junction.^14,15^ CTB therefore represents a potential candidate to target motor neurons *in vivo* for delivery of a cargo.

Indeed, the ability of B_5_ toxoids to undergo retrograde transport in neurons has led to extensive use of CTB-conjugates as neuronal tracers.^16–18^ Tinker *et al*. previously demonstrated that by mimicking the full-length holotoxin, fluorescent CT chimeras could be produced by replacing the toxic A1 domain with a fluorescent protein.^19^ Chen et al. expanded on this work by expressing fusions with heat-labile enterotoxin IIA, and showing that these could be successfully delivered to the cytosol of neurons in mammalian cell culture.^20^ This approach, however, requires expression of A2 fusions with the desired cargo. Successful formation of AB_5_ complexes relies on assembly of the B_5_ pentamer around the A2 α-helix, making co-expression a necessity (Figure 1).^21^ For development of a tool to deliver a payload of any kind, co-expression would ideally be avoided, as not all molecules of interest are amenable to forming fusions, e.g. nucleic acids and antibodies, and separate expression would prevent extended handling of potentially toxic complexes. Identifying a molecule capable of associating with preassembled CTB pentamer in a non-covalent manner could circumvent these problems, since the cargo of choice could be ligated to the CTB-binding component. Each component could then be produced independently and assembled immediately prior to administration.

**Figure 1.**
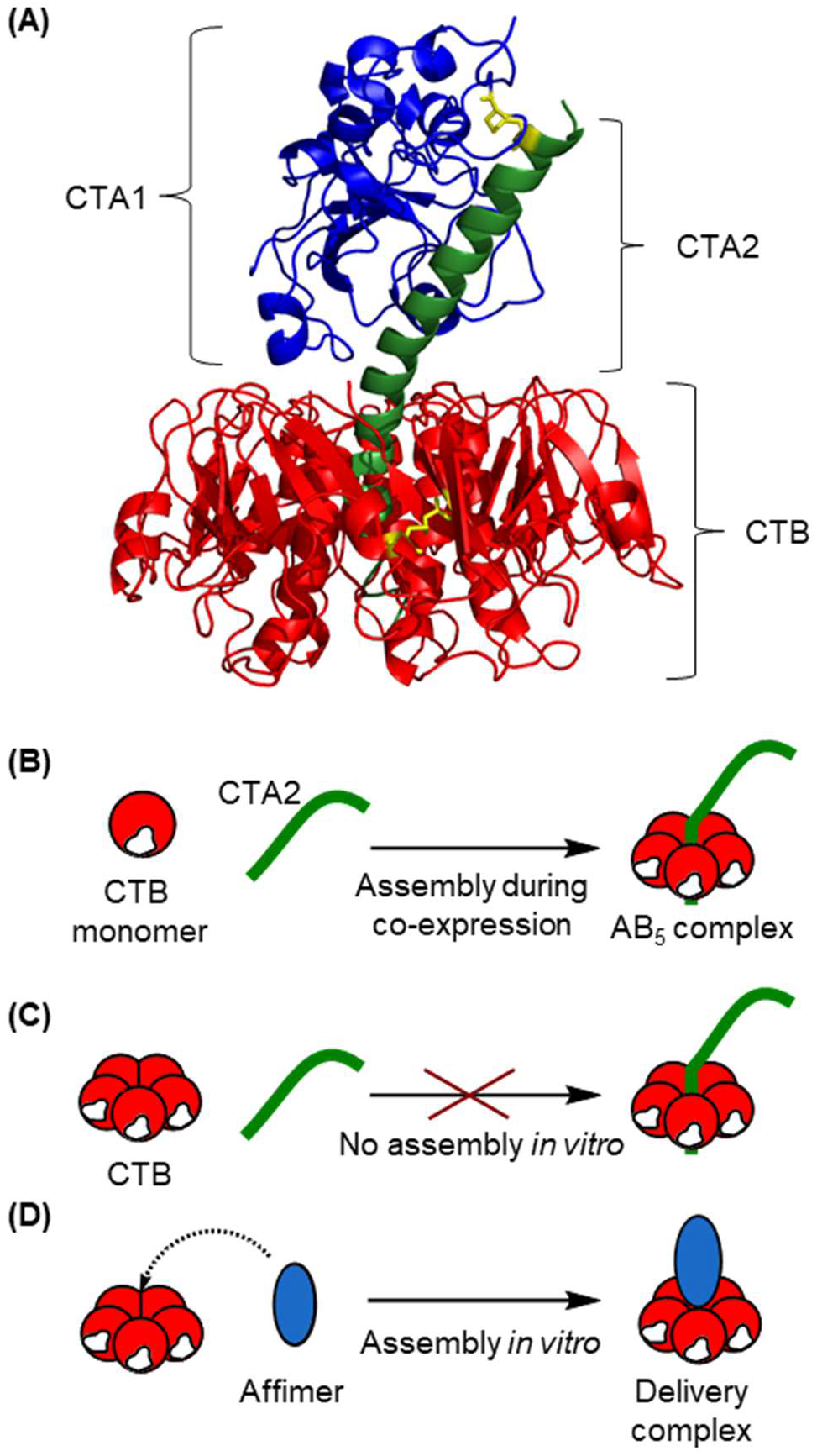
(A) Crystal structure of the AB_5_ cholera toxin showing the CTA1 subunit (green) associated to CTA2 subunit (green) via a disulfide bond (yellow) which in turn is non-covalently associated with the B_5_ subunit CTB (red). (B) Co-expression of A- and B-components leads to assembly of AB_5_ complex *in vivo*, while (C) no spontaneous assembly of AB_5_ complexes occurs upon mixing the B-pentamer with the A2 peptide. (D) The approach described in this paper enabling separate expression of components and simple assembly in *vitro* to form a delivery complex

Antibody mimetics represent a group of engineered protein scaffolds which reproduce the molecular recognition of antibodies while possessing enhanced properties. These improvements include: minimal size, stability, and ease of expression.^22,23^ Specificity for a particular target is obtained by screening diverse libraries using technologies such as phage, mRNA, or ribosome display that allow the binding capability of the isolated protein to be linked back to its coding sequence.^24–26^ This strategy for high-throughput identification of binding proteins has led to the discovery of novel interactions with affinities comparable to those attained with antibodies.^27^ One recent addition to the family of antibody mimetics is the Affimer, a thermostable scaffold based on a phytocystatin consensus sequence, with two variable nine-residue loops responsible for substrate binding.^28,29^ The availability of this scaffold library led us to postulate that we could identify an Affimer to target the non-binding face of CTB to replicate the formation of an AB_5_-type complex in *vitro*, and that this complex would be capable of internalisation into motor neurons and transportation to the cell body in a retrograde fashion to deliver a payload.

Herein we report the first step toward developing this novel technology, in which we use phage display to identify Affimers associating with the non-binding face of CTB. Two binders were identified from an initial screen, termed ACTA-A2 and ACTA-C6, and were shown to form stable complexes with CTB. The technology was then validated in *vivo* by tongue injection in mice followed by detection in neuronal cells of the brain.

## RESULTS AND DISCUSSION

### Identification of CTB-binding Affimers

CTB for us in phage selections and other downstream processes detailed in this paper was expressed in *Vibrio sp*. 60 and subsequently purified directly from the culture supernatant,^30^ as the protein is released from the periplasm following accumulation. Typical approaches to identify target-specific binding proteins by phage display rely on modification of the target to permit immobilisation to a solid support.^31–33^ Recombinant bacteriophage libraries can then be tested against the fixed target in multiple screens to obtain a series of binders. However, modification can be non-specific, resulting in the target being displayed in a variety of orientations. It was critical that Affimer-association did not interfere with the ability of CTB to bind to GM1 as this would impede cellular uptake; thus presentation of CTB in a range of orientations was undesirable. CTB-GM1 complex formation benefits from a particularly high binding affinity (monovalent *K*_d_ = 43 nM) for a protein-carbohydrate interaction,^34,35^ allowing phage display screens to be carried out against unmodified CTB complexed to ganglioside GM1. Adhering ganglioside GM1 to a plate surface ensured that the binding face of CTB was obscured during screening, and that the non-binding face was targeted exclusively. Four consecutive rounds of phage display were conducted in this manner.

Binding of Affimer-displaying phage particles to CTB was confirmed in the first instance by phage ELISA (Figure S1) which led to the identification of fifteen unique clones. In order to prioritize a subset of binders for full characterisation, a plate-based Affimer-lectin binding assay (ALBA) was established. Affimer coding sequences were subcloned from a phage vector into pET11, to add a C-terminal cysteine and His-tag, with the overexpressed protein biotinylated via maleimide modification. Biotinylated Affimer was then evaluated for its ability to bind to GM1-coated plates in the absence or presence of CTB. The amount of Affimer remaining on the plate surface following extensive wash steps was quantified using a streptavidin-HRP conjugate in the presence of the fluorogenic HRP-substrate Amplex^®^ Red and hydrogen peroxide (Figure 2). Affimers A2, B11, C6, and D3 showed the highest binding relative to the control wells and were thus taken forward for further study. These proteins shall now be referred to as Anti-Cholera Toxoid Affimers (ACTAs).

**Figure 2.**
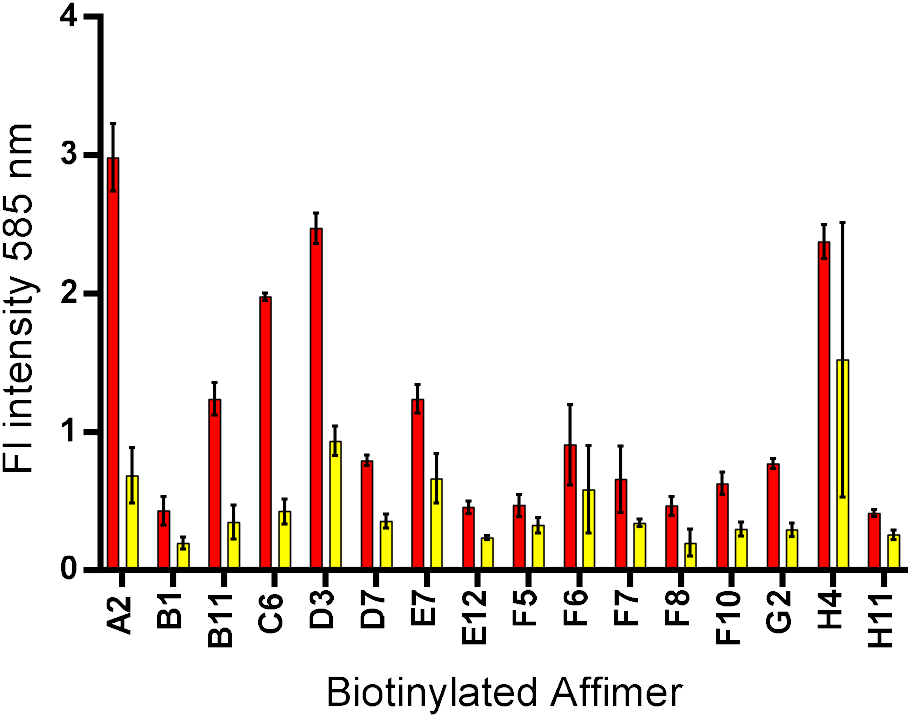
Analysis of CTB-binding Affimers using the Affimer-lectin binding assay (ALBA). Each data point represents the signal produced following incubation of a biotinylated Affimer with (red) or without (yellow) CTB, detected using a Streptavidin-HRP conjugate in the presence of Amplex Red and hydrogen peroxide. The fluorescence of the Amplex Red reaction product resorufin was detected at 585 nm and measured in technical triplicate. Error bars show the standard deviation.

### CTB-binding Affimers Demonstrate Selectivity

In an effort to understand the binding specificity of ACTAs to CTB relative to other similar lectins, the CTB homologue heat-labile enterotoxin B subunit (LTB) was employed. This lectin is 80% identical to CTB, with many of the differentiating residues exposed on the surface of the protein.^36,37^ A similar ALBA was set up, with the five strongest binding proteins to CTB being tested against both CTB and LTB (Figure 3A). ACTA-B11, C6 and D3 appear not to discriminate between the two toxoids. Interestingly, however, ACTA-A2 demonstrated strong selectivity, binding exclusively to CTB. Discrimination between these two highly similar proteins is of particular interest since current commercial antibodies are unable to do so. This assay gave some insight into where ACTA-A2 may be binding, since of the 17 residues that differ between CTB and LTB, only five of these are surface exposed on the non GM1-binding face of CTB. These residues were therefore considered to be the most likely to be forming interactions with the ACTA variable loops (Figure S2).

**Figure 3.**
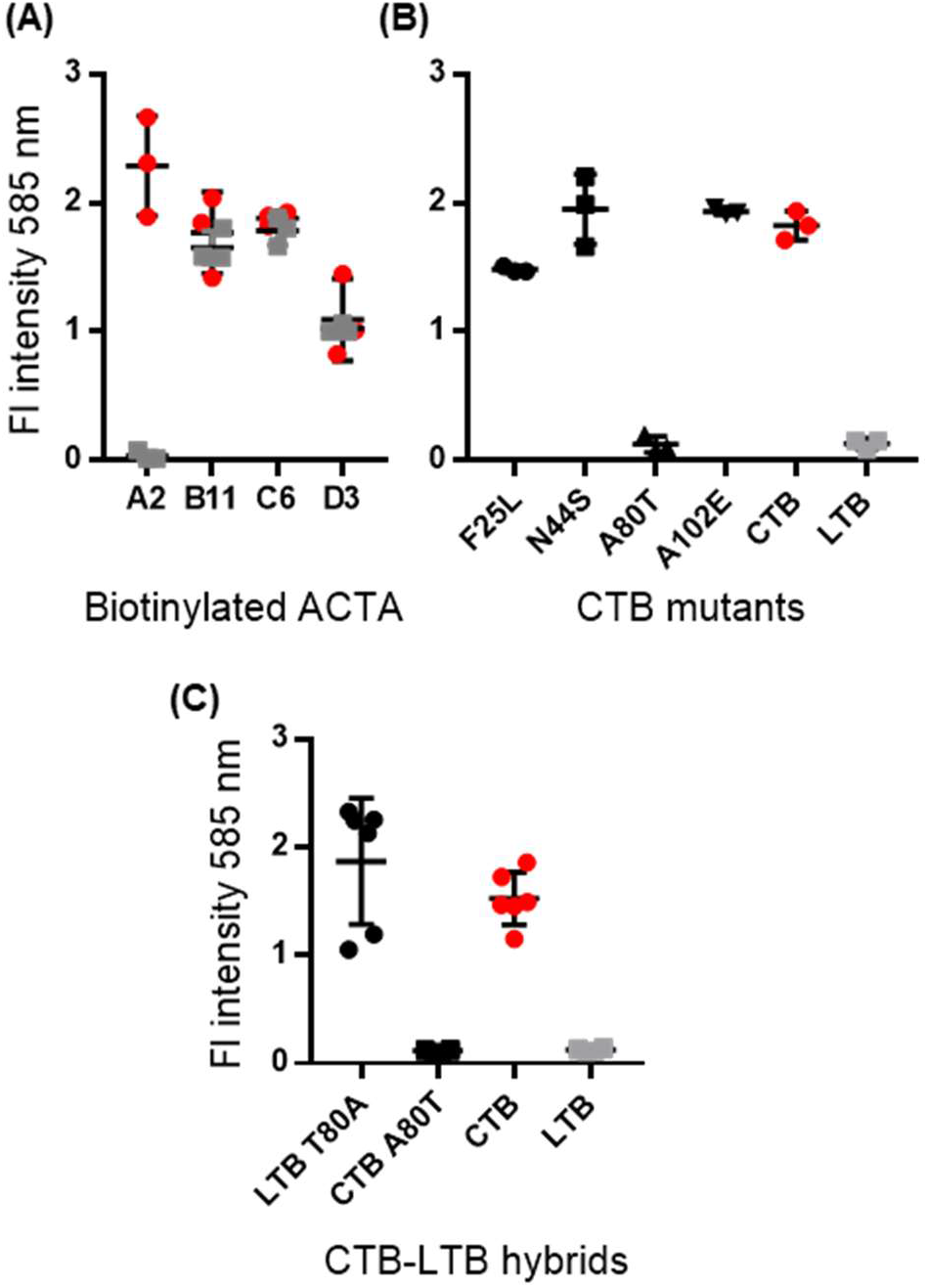
(A) CTB-binding Affimers demonstrate variable selectivity for CTB. Selectivity assays were carried out by incubating GM1 bound CTB (red circles) or GM1 bound LTB (grey squares) with biotinylated ACTA-A2, B11, C6, and D3 in an ALBA assay. (B) ACTA-A2 shows selectivity. Fluorescence detected following testing of ACTA-A2-biotin against a series of five CTB mutants (F25L, N44S, A80T, E83D, A102E, LTB and CTB). (C) A80T facilitates binding of ACTA-A2 to LTB. Fluorescence detected following testing of ACTA-A2-biotin against purified LTB A80T, LTB, or CTB. Signal detected plotted with error bars showing the standard deviation of three technical repeats (A and B) or six technical repeats (C).

To tease out the nature of this interaction, five CTB-LTB hybrids were produced, each carrying a single amino acid mutation corresponding to the five residues postulated to mediate ACTA-A2-CTB selectivity (Figure S3). The hybrid proteins were expressed, and the protein-containing media tested directly in the ALBA with ACTA-A2 biotin (Figure 3B).

Of the hybrids, only one, CTB A80T, no longer bound to ACTA-A2. We therefore generated the corresponding hybrid LTB T80A and tested its interaction with ACTA-A2 biotin in the ALBA (Figure 3C). The mutation resulted in a gain in function for LTB, demonstrating that residue A80 in CTB alone is responsible for the observed selectivity of ACTA-A2 for CTB over LTB. A80 lies in a cleft close to the central cavity at the interface of two CTB protomers (Figure 4) which likely provides a binding pocket for one of the variable regions of ACTA-A2. This site could potentially become unavailable when A80 is mutated to a larger amino acid, such as threonine as found in LTB. Since ACTA-B11, C6 and D3 do not discriminate between the two toxoids, it is unlikely that they bind to the same region as ACTA-A2. Instead a conserved region, such as the pentamer pore, is more likely to be critical for binding. Due to their different binding specificities for CTB and LTB, ACTA-A2 and C6 were selected for further biophysical characterisation.

**Figure 4.**
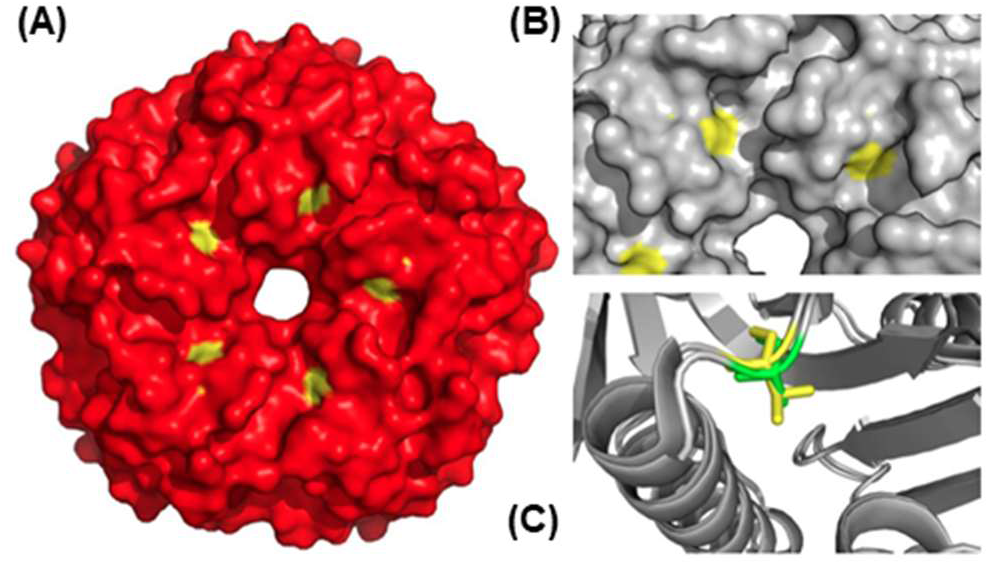
(A) Surface representation of the non-binding face of CTB, with residue A80 coloured in yellow. (B) A close-up image of the hypothesized ACTA-A2 binding groove, showing A80 in yellow. (C) A close-up of an overlay of CTB and LTB with A80 shown in green, and T80 shown in yellow.

Isothermal titration calorimetry (ITC) was applied to measure the binding interaction between each of ACTA-A2 and ACTA-C6 and CTB, with binding constants determined at 37°C to simulate temperature *in vivo*. ITC data is summarized in Table 1. An exothermic heat change was observed for both Affimers, and *K*_d_ values of 1.1 μM and 97 nM were determined for ACTA-A2 and ACTA-C6, respectively. Importantly this revealed that CTB binds to ACTA-C6 10-fold more tightly than to ACTA-A2. ITC also revealed that each ACTA bound to the CTB pentamer as a 1:1 complex. The 1:1 binding for ACTA-A2 to CTB is consistent with it binding to the central pore of the CTB pentamer with one of its loops projecting into the groove distinguished by A80. Interestingly, as ACTA-C6 also displays 1:1 binding, it suggests that this Affimer also binds in the central pore, even though it does not distinguish between A80 and T80 in CTB and LTB.

**Table 1.** Results of ITC experiments

| Affimer | ACTA-A2-CTB | ACTA-C6-CTB |
| --- | --- | --- |
| Temp (K) | 308 | 308 |
| $\Delta H^\circ$ (kcal/mol) | $-23.0 \pm 0.87$ | $-7.33 \pm 0.14$ |
| $\Delta G^\circ$ (kcal/mol) | $-8.40 \pm 1.65$ | $-9.95 \pm 0.19$ |
| $K_d$ (nM) | $1130 \pm 210$ | $97 \pm 30$ |
| $T\Delta S^\circ$ (kcal/mol) | $-14.64 \pm 2.52$ | $2.62 \pm 0.33$ |
| $n^a$ | $1.15 \pm 0.03$ | $1.05 \pm 0.01$ |
| $N^b$ | 3 | 2 |

<sup>a</sup> binding stoichiometry; <sup>b</sup> number of replicates

### Protein delivery in cultured cells

We next assessed the ability of these novel complexes to be taken up into mammalian cells prior to validation *in vivo*. As a first proof of concept, ACTA proteins and CTB were labelled with Alexa Fluor 555 and 488 respectively. Sitespecific routes to labelling were carried out to ensure that complex formation would not be impeded by chemical modification. Affimers harbouring a C-terminal cysteine were modified with maleimide fluorophores, while CTB was oxidized with sodium periodate and then modified at its N-termini with aminooxy fluorophores.^38^ ACTA:CTB complexes were produced by combining the toxoid with the Affimers in a 1:1 ratio and subsequently purified by gel filtration (Figure S4). A single peak containing ACTA-555 co-eluted with CTB-488 was collected, and the protein was administered to Vero cells, a cell-line derived from monkey epithelial cells expressing ganglioside GM1. The presence of labelled proteins internalized within the cells was readily detected at concentrations as low as 175 nM and 15 nM for ACTA-A2/CTB and ACTA-C6/CTB respectively (Figure 5 and Figure S5). The difference in affinity between ACTA-A2 and ACTA-C6 for CTB was thus reflected in the minimal concentration threshold required for detection of labelled complex in cell culture.

**Figure 5.**
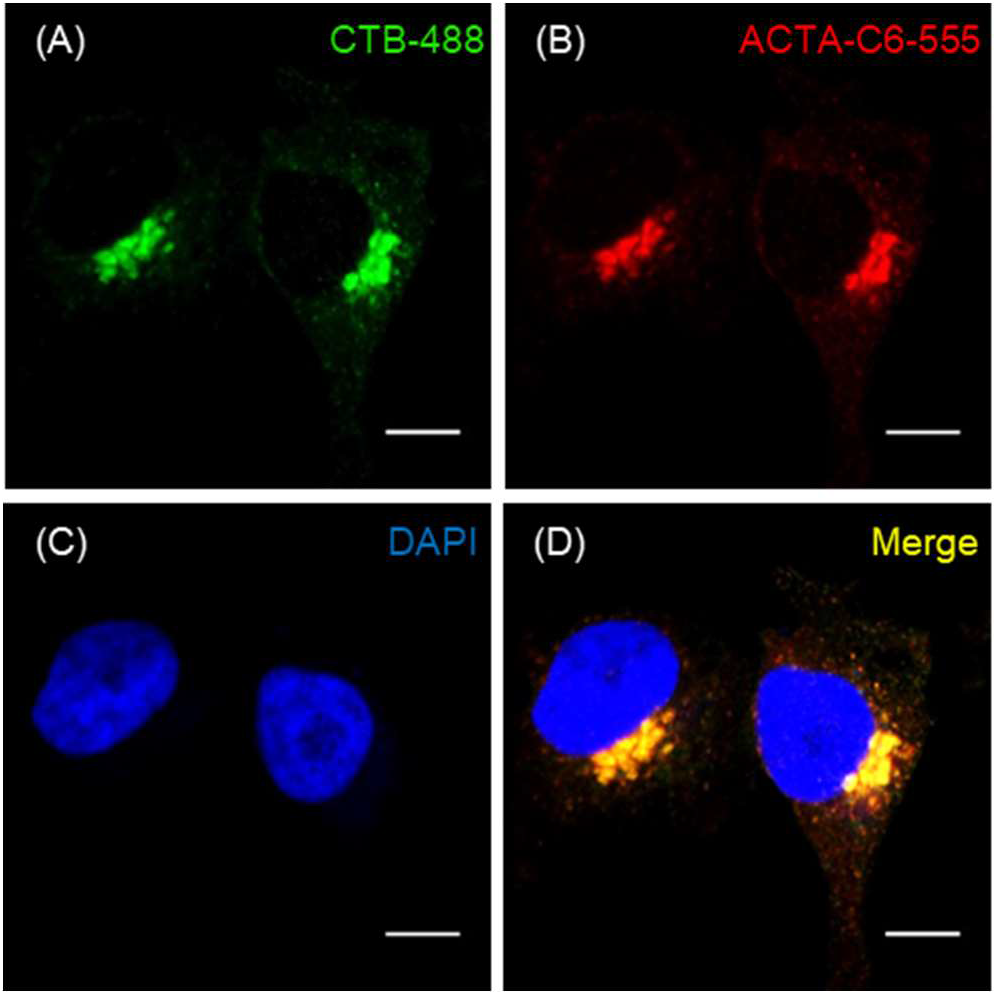
Vero cells treated with CTB-488/ACTA-C6-555 complex to a final concentration of 15 nM and stained with DAPI nuclear stain. Scale bars are 10 μm. (A) CTB-488 fluorescence, (B) ACTA-C6-555 fluorescence, (C) DAPI nuclear stain blue fluorescence, (D) merged image of 488, 555, 405 nm wavelengths.

Analysis of the cell images reveals that CTB and ACTAs clearly co-localize and appear concentrated in discrete puncta in an area directly adjacent to the nucleus that is consistent with the morphology of the Golgi apparatus. CTB is known to enter recycling endosomes, and be transported to the trans-Golgi network from the cell surface, resulting in accumulation of CTB in the Golgi apparatus.^39^ Preliminary analysis of complex localisation therefore appears to agree with previous observations and indicate that ACTA complexes do not alter the endocytic pathway of CTB.

### Delivery of Affimers to motor neurons

Numerous studies report the ability of CTB to be endocytosed into neurons when delivered to the CNS.^40,41^ Neuronal tracing in this capacity has been achieved by intraperitoneal and intravenous injections for broad dispersal of CTB into the neurons of the spinal cord and brain.^42^ A more targeted delivery approach that is useful for testing putative reagents, with labelling specifically of the neurons of the hypoglossal nucleus in the brainstem, has been described by intramuscular administration of protein into the tongue,^43,44^ therefore, injections were carried out paralingually. C57BL/6 mice were treated with either 25 μg unlabelled CTB complexed with one of the two fluorescently-labelled Affimers (ACTA-A2-555 or ACTA-C6-555), or with one of the three individual components (5 μg ACTA or 20 μg CTB). After 24 hours, cross-sections of the brainstem were prepared and stained with a primary-secondary antibody pair to detect endocytosed CTB. Slides were then viewed by confocal microscopy (Figure 6).

**Figure 6.**
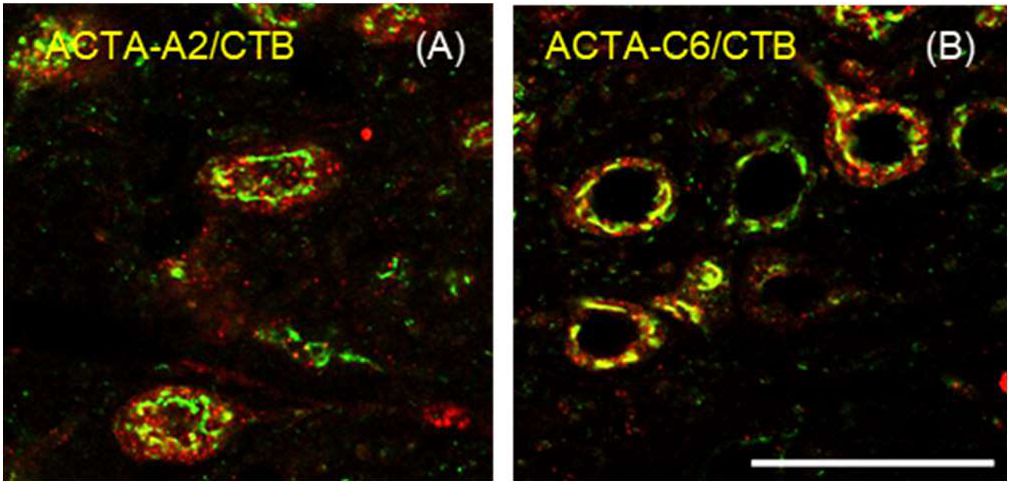
Cells of the hypoglossal nucleus in brainstem crosssections for mice paralingually injected with ACTA-A2-555/CTB (left, image (A)) or ACTA-C6-555/CTB (right, image (B)), showing ACTA fluorescence (555 nm, red) overlaid with CTB immunolabelling (488 nm, green). Scale bar = 50 μm.

CTB staining and ACTA fluorescence were both observed in the brainstems of mice treated with the complex, but not in mice treated with the fluorescent ACTA alone (Figure S6) demonstrating that ACTA internalisation was dependent on CTB. In a similar manner to earlier experiments carried out in Vero cells, localisation could broadly be assigned to a region peripheral to the motor neuron nucleus. Detection of ACTA-CTB at the nucleus indicates complete retrograde trafficking of the complex from the neuromuscular junction to the cell body, contained within the brainstem.

The two ACTA-CTB complexes showed different localisation patterns. After 24 hours, ACTA-A2 is observed in small punctate spots throughout the cell, while the fluorescence for ACTA-C6 is peripheral to the nucleus as observed in preliminary cell experiments (see Figure 5/Figure S5). These halos of ACTA-C6-555 fluorescence overlay with the detected CTB, suggesting that the higher affinity ACTA-C6 is still bound to its target after internalisation and trafficking to the cell body. Conversely ACTA-A2-555 fluorescence overlaid poorly with CTB, suggesting that the complex dissociates after internalisation and the stable Affimer is then being sequestered inside endosomes for lysosomal degradation.

A further experiment to establish the longevity of the complex *in vivo* was carried out by administering the complex paralingually and sacrificing the mice after one week. Few ACTA-A2-555 positive cells were detected, with nearly all remaining fluorescence appearing clustered in punctate spots (Figure 7). ACTA-C6-555, however, was readily detected in the hypoglossal nucleus, with much of ACTA-C6-555 fluorescence overlaying with immunostained CTB, indicating a longer-lived half-life for the ACTA-C6/CTB complex. We propose that the lower affinity ACTA-A2 dissociates from CTB rapidly, resulting in a reduced half-life *in vivo* compared to ACTA-C6, which remains bound for longer, leading to an extended half-life (Figure 8). Of note is that Affimers are derived from phytocystatins, a group of lysosomal protease inhibitors found in plants. Lysosomal degradation of these proteins might therefore be surprising, although the ability of phytocystatins to survive degradation in a mouse lysosome is unknown. It is possible that ACTA-A2 persists in the motor neuron, albeit without the fluorescent tag.

**Figure 7.**
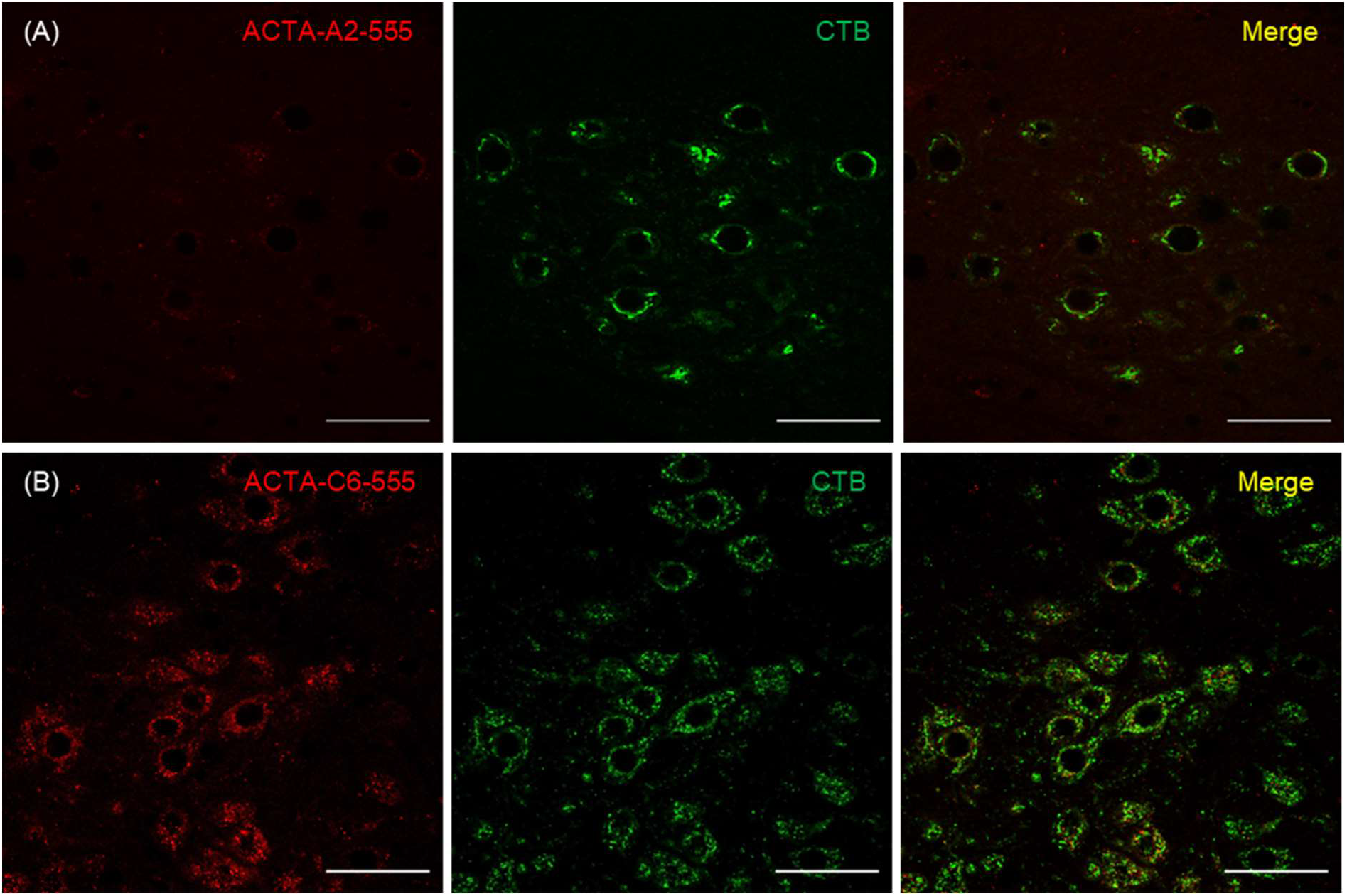
Hypoglossal nucleus in cross-section of brainstems of mice injected paralingually with (A) ACTA-A2-555 + CTB and (B) ACTA-C6-555 + CTB. Mice were sacrificed one week following injections, with CTB detected by immunolabelling. Scale bars = 50 μm.

**Figure 8.**
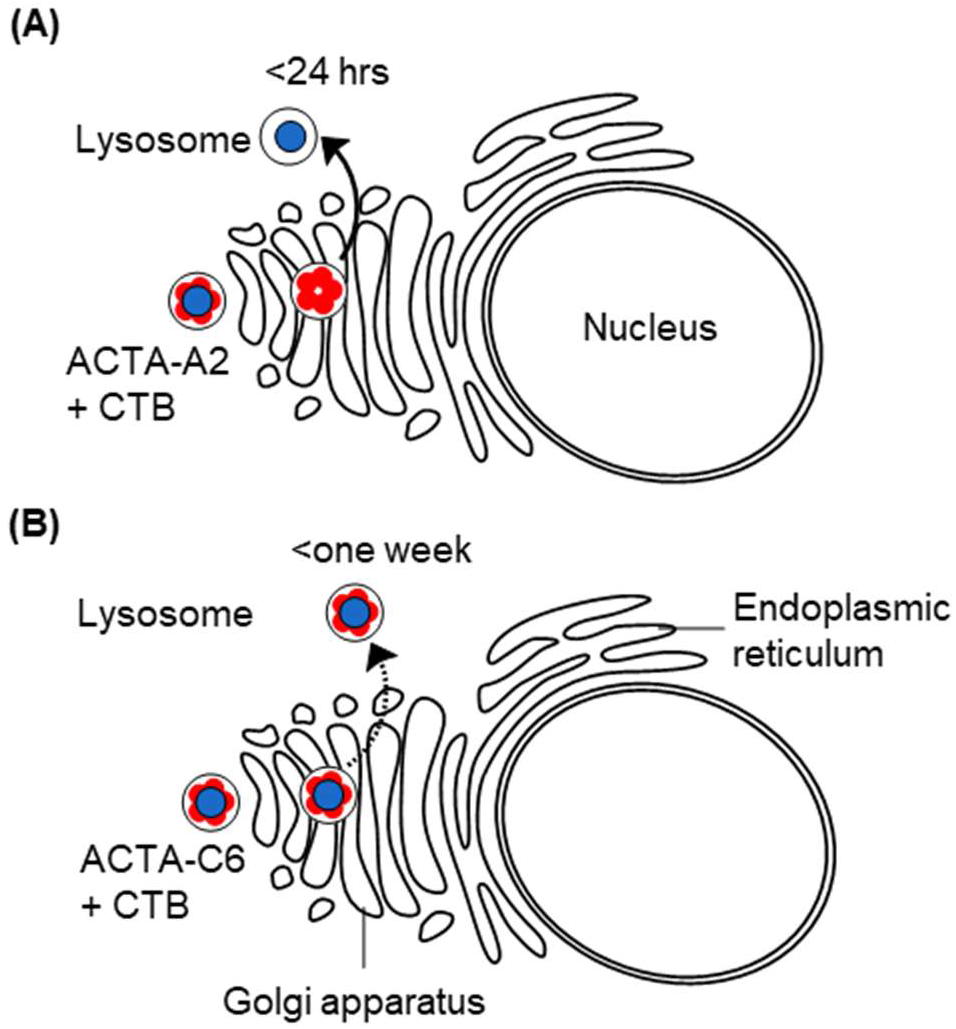
Hypothesis of the fate of the ACTA-CTB complex once internalized into motor neurones. (Top) ACTA-A2-CTB complex is trafficked to the Golgi apparatus in the cell body. Once inside the neuron, ACTA-A2 dissociates from CTB, and is rapidly targeted to lysosomes for degradation. (Bottom) ACTA-C6-CTB complex is trafficked to the Golgi apparatus. The complex remains associated and is eventually targeted for lysosomal degradation.

## CONCLUSIONS

CTB has been extensively studied as a tool for neuronal tracing and protein delivery. In this manuscript we expand on the capabilities of CTB by proposing a novel method for carrying macromolecular cargo into the brainstem of a mouse as a first step to developing a motor neuron-selective protein delivery system. By exploiting the GM1-binding properties of the cholera toxin B-subunit, two anti-cholera toxoid Affimers were identified through phage display that target the face distal to the GM1-binding sites of CTB. Using an ELISA-type assay, selectivity for CTB over LTB was demonstrated for ACTA-A2, but not for ACTA-C6. Fluorescently labelled ACTA-CTB assemblies were injected intramuscularly into the tongues of mice and were readily detected in the motor neurons of the hypoglossal nucleus. Intracellular distribution *in vivo* was noticeably different between ACTA-A2/CTB and ACTA-C6/CTB treated mice, with ACTA-C6/CTB remaining largely associated, whilst ACTA-A2/CTB appeared to have fully disassembled. After 24 hours ACTA-A2 was detectable in discrete vesicles but had significantly reduced after one week. The weaker interaction of the ACTA-A2/CTB complex was hypothesized to be facilitating disassembly *in vivo*. A lower affinity may be preferable for release of cargo following internalisation and may be necessary to achieve dispersion throughout the neurons of the body. As a proof of concept, we have shown that CTB can be used as a vehicle to deliver a piggybacking cargo into motor neurons. Beyond the use of the CTB:Affimer approach described here, this approach could be applied to any antibody-mimetic and lectin pair to develop a modular protein assembly for intracellular delivery of macromolecular cargo. This strategy therefore has the potential to be adapted to delivery of molecular cargoes to a wide variety of cell types beyond those which interact with CTB.

## METHODS

### Affimer Expression

See Tiede *et al*.^28^ for detailed methods. Briefly, an overnight culture of BL21(DE3) cells transformed with pET11-Affimer was used to inoculate 400 mL LB-ampicillin medium. Cultures were induced with 0.5 mM IPTG at OD 0.6 and incubated at 30 °C for at least six hours before chemical cell lysis. Clarified supernatant was purified by immobilised nickel affinity chromatography, followed by size exclusion chromatography.

### CTB Expression from *Vibrio sp. 60*

El Tor CTB was expressed from Vibrio sp. 60 transformed with pATA13 as described previously.^30^ In brief, an overnight culture incubated at 30 °C was used to inoculate 1 L high-salt LB medium (2 % NaCl). At OD 0.6 protein expression was induced with IPTG (0.5 mM) and the cells were incubated at 30 °C for 24 hours. Cells were pelleted by centrifugation and protein was purified directly from the medium by ammonium sulfate precipitation at a final concentration of 57% w/v ammonium sulfate. The saturated solution was then stirred for two hours at room temperature then centrifuged at 18,000 x*g* for 30 minutes. The supernatant was discarded and the pellet was resuspended in 20 mL PBS. A second centrifugation was carried out to remove insoluble material. Protein was then purified using a lactose affinity column at 4°C, and eluted in a lactose-rich buffer (50 mM phosphate pH 7.4, 300 mM NaCl, 300 mM lactose).

### Phage Display

Phage selections and phage ELISA were carried out as detailed in Tiede *et al*.^28^ GM1-coated plates were prepared by adsorption of 100 μl 1.3 μM solution of GM1 ganglioside in methanol to Nunc MaxiSorp flat-bottom 96 well plates. The methanol was allowed to evaporate, and the GM1 plates were blocked overnight by incubation at 37°C with a casein-based blocking buffer. Plates were washed once with 300 μl PBST on a plate washer immediately before use. Three rounds of panning were carried out on GM1-coated wells treated with 50 μl 1 μM solution of CTB captured for one hour at RT then washed three times with PBS + 0.05 % TWEEN 20 (PBS-T).

### Affimer-lectin binding assay

Affimers were biotinylated using a 20-fold molar excess of biotin-PEG2-maleimide in 50 mM phosphate pH 7.4, 300 mM NaCl supplemented with 1 mM TCEP. Reactions were incubated overnight at RT then dialysed into PBS. Greiner Bio-one high binding black 96 well plates were adsorbed with ganglioside GM1 as described in the previous section. 50 μl lectin (CTB/LTB or similar) (1 μM) was transferred to the appropriate wells, and left to bind for one hour at RT. Plates were washed once with PBS-T, and 100 μl biotinylated Affimer (250 nM) was added to each well. Plates were washed once after Affimer binding at RT for one hour. 50 μl streptavidin-HRP complex (Ultra Streptavidin-HRP, Life Technologies) diluted at 1:1000 in PBS was added to the wells of the plate. The one hour incubation was repeated and the plate was washed six times. A solution of Amplex Red and hydrogen peroxide in PBS, both at a final concentration of 5 μM, was made up and 50 μl of this substrate was added per well. Plates were loaded into a plate reader for fluorescence detection at 585 nm.

### Labelling Affimers with Alexa Fluor 555 C2 maleimide

400 μl Affimer C-term Cys (150 μM) in 50 mM phosphate, 150 mM NaCl, 0.5 mM TCEP, pH 7.2 was combined with 0.94 mg Alexa Fluor 555 C2 maleimide (equal to five molar equivalents). Samples were incubated at room temperature for four hours in foil, before purification using a PD-10 desalting column, concentration to 250 μl, and purification by gel filtration using an analytical Superdex 75 increase column.

### Labelling CTB with aminooxy Alexa Fluor 488

CTB was labelled with aminooxy Alexa Fluor 488 by oxidising the N-terminal threonine, and combining the subsequent oxidised CTB with two molar equivalents of the lyophilised fluorophore. CTB (500 μM [protomer concentration], in PBS), L-methionine (10 equivalents), and NaIO4 (5 equivalents) in phosphate buffer, were incubated together at room temperature out of light for 45 minutes, before buffer exchanging into PBS (**pH 6.8**) using a PD10 desalting column. The eluate was concentrated to 0.5 mL (~500 μM), and added to 0.4 mg lyophilised fluorophore (2 molar equivalents) and 4.5 μl aniline (neat). To reduce the number of fluorophores per pentamer, the amount of fluorophore can be reduced to 0.4 molar equivalents. After a 20 hour incubation, the protein was purified using a G25 desalting column, concentrated to 0.5 mL, and purified by gel filtration.

### Isothermal Titration Calorimetry

All ITC experiments were performed using a MicroCal ITC_200_. An initial sacrificial titration of 0.5 μl was required, before 19 x 2 μl titrations were carried out, each spaced out over 4 seconds, with a 120 second delay in between each injection. Experiments were initially carried out at 100 μM CTB, titrated into 10 μM Affimer. These concentrations were then varied, if necessary, to obtain a sigmoidal shape of curve. For each set of concentrations, a CTB titration into buffer was carried out to obtain a measure of the heat of dilution of CTB into buffer. These data could then be subtracted from the data obtained from ligand into receptor titrations to bring the baseline closer to zero, thereby facilitating curve fitting. Data were analysed and fitted with a ‘one site bind’ model using Origin Microcal software.

### Cell Preparation and Analysis

Seeded cells (1.2 × 10^5^ cells per well) in 12-well plates to be treated with protein were washed, by replacing the Cell Culture Media with 1 mL fresh media. Protein solution was then added directly to the media, and the cells were incubated for 4-6 hours at 37 °C, 5% CO_2_ prior to cell fixation. Growth medium was removed, and the cells were washed three times with PBS. 0.5 mL Cell Cleansing Buffer was added, and incubated at 4 °C for 6 minutes. The cleansed cells were then washed an additional three times, before addition of 4% v/v paraformaldehyde (PFA) in PBS for 30 minutes at room temperature in a fume hood. The PFA was removed, and the cells were washed three times with PBS. Cells were then permeabilised with 0.1% Triton X-100 in PBS for one hour at room temperature in preparation for immunohistochemistry.

### Animals

All animal experimentation was carried out under a Home Office License, and in accordance with the regulations of the UK Animals, Scientific Procedures, Act 1986. Experiments were performed on young male and female C75BL/6 adult mice, bred in house.

The tongue was targeted for intramuscular injections of protein and protein complexes, since it provided a route to deliver material into the motor neurones of the brainstem. CTB-ACTA complexes were prepared at 200 μM then injected via paralingual administration of 2 μl of protein material. Mice were sacrificed after 24 – 96 hours. Brainstems were removed, fixed in 4% PFA for 24 hours at 4°C, then prepared for sectioning. All sections were taken transversely on a VT1000S vibrating microtome at 50 μm thickness. The sectioned slices were collected and transferred into PBS in a 24-well plate. Tissue sections and cultured cells were permeabilised and treated with rabbit anti-CTB diluted 1:1000 in PBS and incubated overnight at 4 °C. A 1:1000 dilution of a fluorescent secondary antibody in PBS was prepared and added to the sample for one hour, then removed and washed three times with PBS. The sample was then air-dried onto a glass microscope slide before addition of 10 μl of Vectashield Mounting Medium and protection of the sample with a cover slip ready for visualisation.

### Confocal Microscopy

For confocal analysis of samples, an Axio Imager Z2 LSM880 upright confocal microscope (Zeiss) equipped with a 405 nm diode laser, Argon 458/488/514 nm, DPSS 561 nm and HeNe 633 nm lasers and a GaAsP detector. Images were captured using the Zeiss LSM Image browser software, and processed using Zen lite 2.3 software. For wide-field analysis of samples, a Nikon Eclipse E600 microscope equipped with epifluoresence and Q-Imaging Micropublishing 5.0 camera was used. Images were captured using AcQuis image capture software.

## ASSOCIATED CONTENT

Detailed experimental procedures and supplementary data presented in Supporting Information.

## AUTHOR INFORMATION

### Author Contributions

All authors have given approval to the final version of the manuscript.

## ACKNOWLEDGMENT

This work was supported by the Wellcome Trust and the Biotechnology and Biological Sciences Research Council – see funding sources.

## ABBREVIATIONS

ACTA: anti-cholera toxoid Affimer
ALS: amyotrophic lateral sclerosis
CNS: central nervous system
CPP: cell-penetrating petides
CTB: cholera toxin B-subunit
HRP: horse-radish peroxidase
ELISA: enzyme-linked immunosorbent assay
ALBA: Affimer-lectin binding assay
LTB: heat-labile enterotoxin B subunit
ITC: isothermal titration calorimetry.

