## Supplementary Information for "Piggybacking on the Cholera Toxin: Identification of a Toxoid-binding Protein as an Approach for Targeted Delivery of Proteins to Motor Neurons"

SUPPORTING INFORMATION

#### Supplementary figures

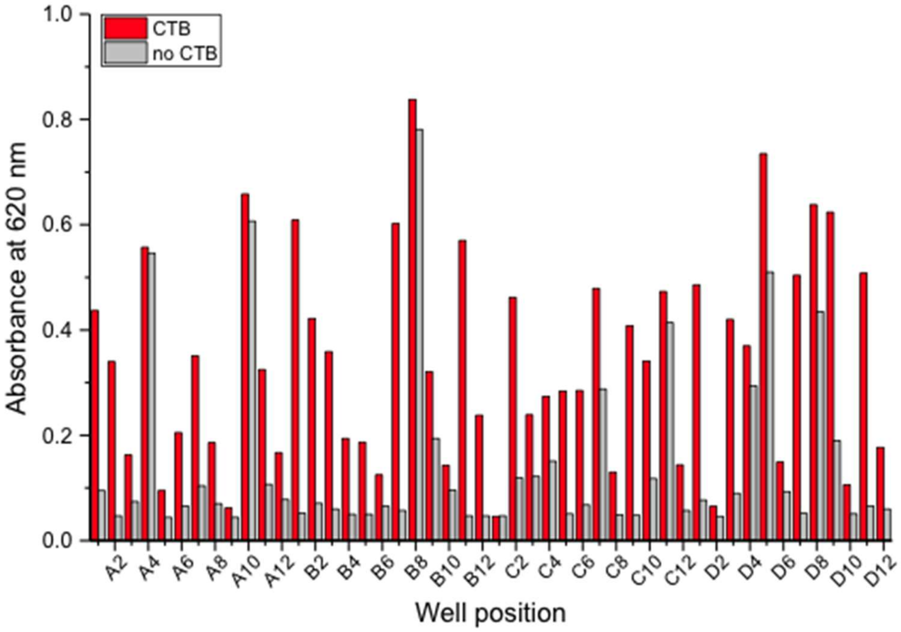

*Supplementary Figure 1. Analysis by phage ELISA of 48 Affimers isolated following panning against wild-type CTB. A single species of Affimer is tested against either CTB bound to GM1 (red bars), or GM1 alone (grey bars). Absorbance at 620 nm is plotted in a bar chart against the well position of the tested Affimer to facilitate identification.*

| | Variable loop 1 | | | | | | | | | Variable loop 2 | | | | | | | | |
|---|---|---|---|---|---|---|---|---|---|---|---|---|---|---|---|---|---|---|
| **A2** | Q | H | E | R | S | H | W | V | D | H | N | Q | F | F | D | Y | F | I |
| **B1** | P | P | D | S | T | E | Q | Q | R | K | W | P | G | K | F | N | K | Y |
| **B3** | E | F | S | S | S | R | R | V | K | G | L | S | T | I | G | K | I | L |
| **C4** | V | D | Q | K | P | P | A | R | M | K | N | F | W | F | P | S | Q | N |
| **C6** | M | D | L | N | A | G | L | P | R | Q | G | L | K | K | L | K | F | T |

*Supplementary Figure 2. Primary sequences of the variable loops of ACTAs identified following selections against CTB. Colour scheme corresponds to RasMol colour grouping by traditional amino acid properties.*

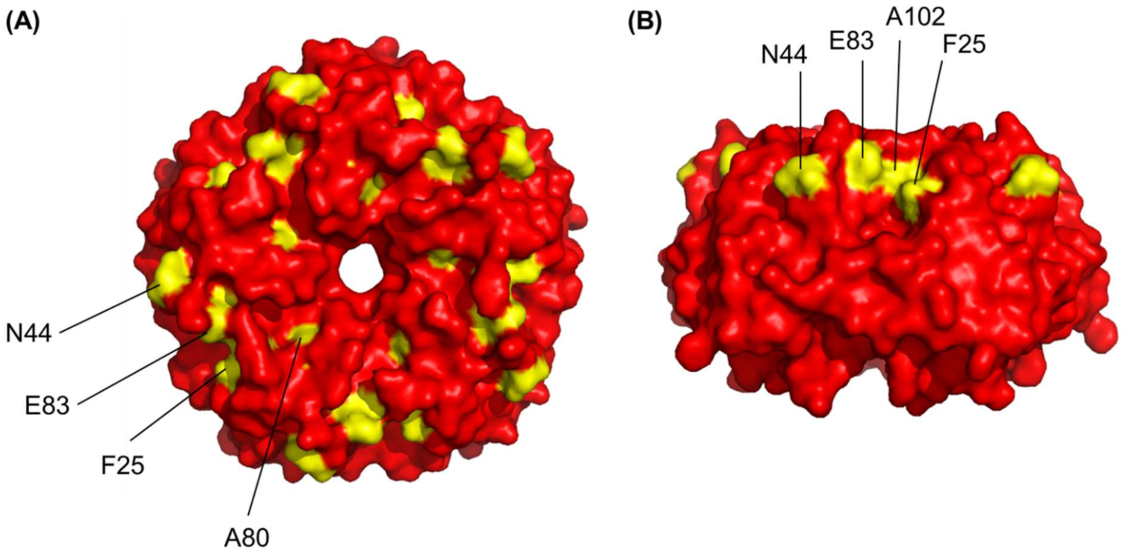

*Supplementary Figure 3. (A) A surface representation of the top face of CTB. There are five surface exposed residues per protomer that differ between the top faces of CTB and LTB. These residues have been highlighted in yellow. (B) A side on view of CTB with surface exposed residues differing to LTB coloured in yellow.*

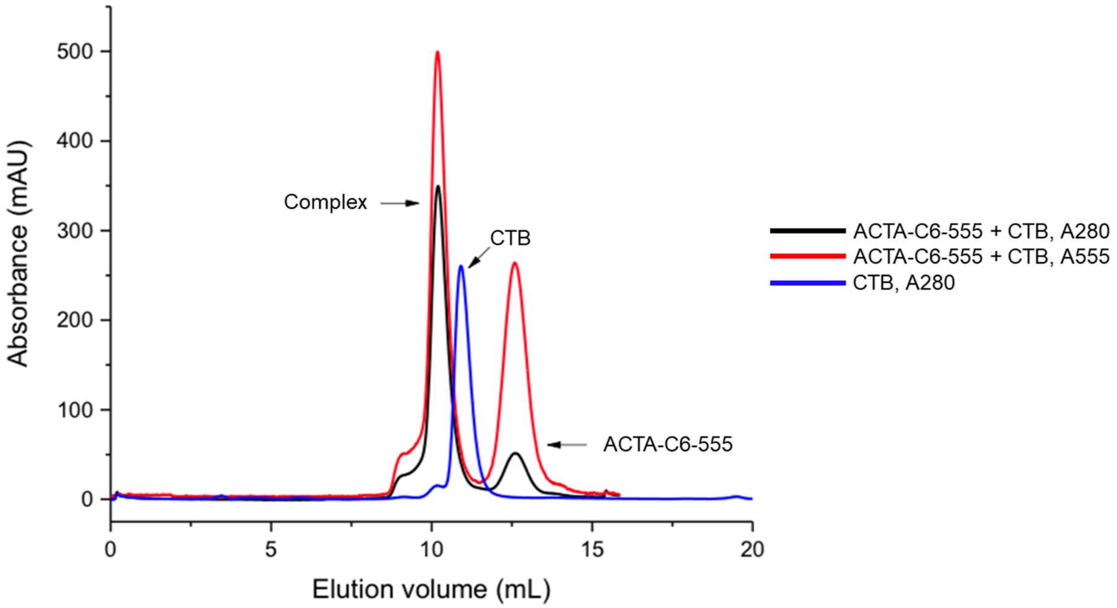

*Supplementary Figure 4. Gel filtration chromatogram generated following the purification of C6-555:CTB complex using a Superdex 75 10/300 GL gel filtration column, with detection at 280 nm (black) and 555 nm (red) absorbance. Arrows denote the complex formed and free unbound C6-555 (Affimer C6 labelled with Alexa Fluor 555). A chromatogram of CTB (blue) has been overlaid as a reference to show the shift in elution volume of the complex.*

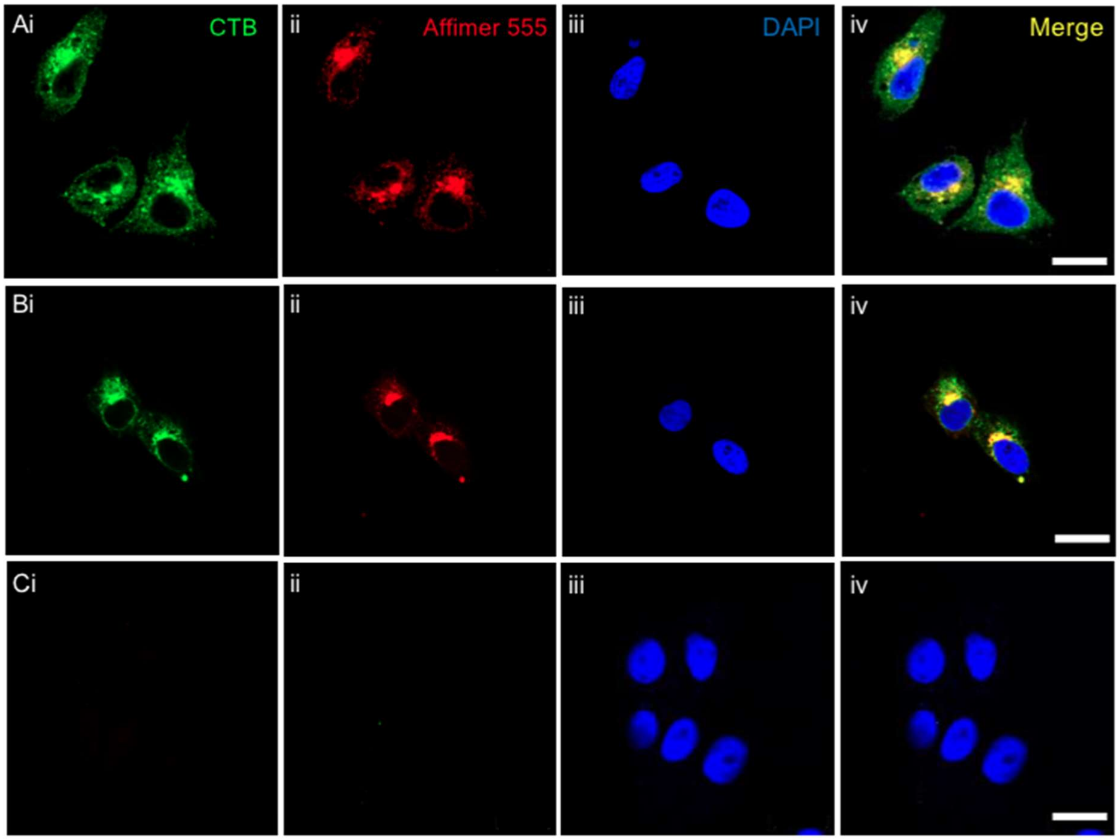

*Supplementary Figure 5. Vero cells incubated with (A) C6-555 + CTB, (B) A2-555 + CTB, (C) C6-555 only, at 750 nM, fixed after six hours, and immunolabelled for CTB (panel i) and DAPI (panel iii). Scale bars are 20 µm.*

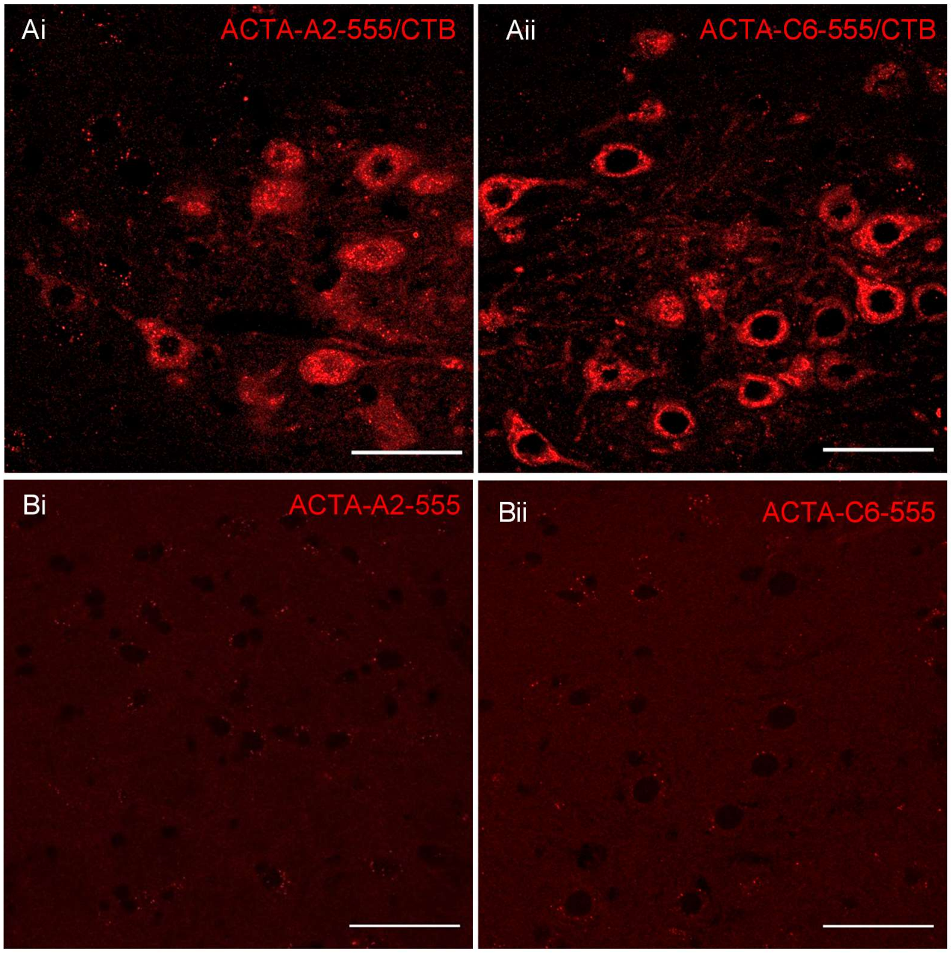

*Supplementary Figure 6. Magnified cross-sections of the brainstem focusing on the hypoglossal nucleus showing (A) mice injected paralingually with ACTA-555/CTB, ACTA-A2 (i), ACTA-C6 (ii) (B) mice injected paralingually with ACTA-555 alone, ACTA-A2 (i), ACTA-C6 (ii). Scale bars are 50 µm.*

### 1 Materials and Methods

#### 1.1 Standard Buffer Solutions

All common buffers were adjusted to pH 7.4, unless otherwise stated, and made up with 15 MΩ water to the required volume.

**Phosphate Buffer:** 50 mM $NaH_2PO_4$, 150 mM NaCl

**Phosphate Buffered Saline (PBS):** 10 mM $Na_2HPO_4$, 1.8 mM $KH_2PO_4$, 137 mM NaCl, 2.7 mM KCl

**PBS-T:** 50 mM $NaH_2PO_4$, 150 mM NaCl, 0.05% TWEEN20

**Tris Buffered Saline 1X (TBS):** 50 mM Tris-HCl, 150 mM NaCl

**Lysis Buffer:** 50 mM $NaH_2PO_4$, 300 mM NaCl, 1 µg $mL^{-1}$ DNAse I, 1x Vol Halt Protease Inhibitor

**Lactose Elution Buffer:** 50 mM $NaH_2PO_4$, 300 mM NaCl, 300 mM Lactose

**Ni-NTA Wash Buffer:** 20 mM Imidazole, made up to desired volume with lysis buffer

**Ni-NTA Elution Buffer:** 500 mM Imidazole, made up to desired volume with lysis buffer

**Blocking Buffer:** 20% v/v 10× Casein Blocking Buffer in *PBS-T*

**Cell Cleansing Buffer:** 0.2 M acetic acid, 0.5 M NaCl, pH 2.8

**Solubilisation Buffer:** 10 mM $Na_2HPO_4$, 1.8 mM $KH_2PO_4$, 137 mM NaCl, 2.7 mM KCl, 0.1% Triton-X 100

#### 1.2 Media

**Miller's Lysogeny Broth (LB) Medium:** 1.0% w/v Tryptone 0.5% w/v Yeast Extract 1.0% w/v NaCl

**LB Agar Antibiotic Plates:** 1.5% w/v Agarose, 2.5% w/v LB Medium**.** The agar was supplemented with a final concentration of 100 µg/ml ampicillin or 50 µg/mL kanamycin to generate the selective medium.

**2x Tryptone/Yeast Media (2TY):** 1.6% w/v Tryptone, 1.0% w/v Yeast Extract, 0.5% w/v NaCl

**High Salt LB Growth Media:** 1.0% w/v Tryptone, 0.5% w/v Yeast Extract, 2.0% w/v NaCl

**Cell Culture Media:** DMEM/F-12 GlutaMAX, 10% v/v foetal bovine serum, 5% v/v Penicillin-Streptomycin (Sigma)

##### 1.2.1 Polymerase Chain Reaction

Standard polymerase chain reaction (PCR) was used to amplify DNA for gel extraction, or for product analysis, to ensure that all primers resulted in a single, specific DNA fragment being produced. In all cases, New England Biolabs (NEB) enzymes and buffers were used. Standard PCR was set up in 50 µl reactions on ice, and contained the following:

1× Phusion High Fidelity Polymerase buffer (NEB)

500 nM Forward primer

500 nM Reverse primer

400 µM dNTP

1-25 ng DNA template

1 unit Phusion High Fidelity Polymerase (NEB)

This solution was made up to 50 µl with sterile $H_2O$, and the following cycling conditions were used:

| | | |
|---|---|---|
| 95 °C | 5 minutes | |
| 95 °C | 20 seconds | 30 Cycles |
| $T_m$ | 20 seconds | 30 Cycles |
| 72 °C | 45 seconds | 30 Cycles |
| 72 °C | 5 minutes | |
| 4 °C | Hold | |

An extension time of 45 seconds was suitable for amplification of DNA segments less than 1 kb in size. A further 20 seconds was added for every additional kb of DNA to be amplified.

PCR products were purified using Qiagen QIAquick PCR Purification Kits, used as per the manufacturer's instructions.

##### 1.2.2 Site Directed Mutagenesis

Quikchange site-directed mutagenesis was carried out to change nucleotides within a plasmid in a site-specific manner. The process relies on primers with a high degree of complementarity flanking the region that is to be altered, and hence amplifying the plasmid whilst introducing the desired mutation. DpnI, an enzyme that recognises and cleaves methylated DNA, is added to digest the wild-type plasmid, leaving exclusively the amplified product for transformation. All primers were designed using *Agilent Technologies QuikChange Primer Design* program.[1] Polymerase chain reactions were set up as follows:

1x KOD Hot Start Polymerase buffer (VWR)

140 nM Forward primer

140 nM Reverse primer

1.5 mM $MgSO_4$

200 µM dNTP

50 ng DNA template

1 unit KOD Hot Start Polymerase (VWR)

dd$H_2O$ to 50 µl

Cycling conditions:

| | | |
|---|---|---|
| 95 °C | 2 minutes | |
| 95 °C | 20 seconds | 20 cycles |
| 55 °C | 20 seconds | 20 cycles |
| 70 °C | 2 min 30 sec | 20 cycles |
| 72 °C | 10 minutes | |
| 4 °C | Hold | |

The PCR product was combined with 1 µl DpnI, and incubated at 37 °C for two hours, before transformation into *E. coli* XL10 cells.

##### *1.2.3* Heat-shock Transformation of XL10/BL21 Gold *Escherichia coli*

Chemically competent *E. coli* cells were transformed via heat-shock. *E. coli* XL10-Gold cells were used for plasmid production, whilst BL21 Gold cells were employed for protein expression.

10 µl of cells were combined with 1 µl of plasmid DNA in sterile $H_2O$, and kept on ice for 30 minutes. Following this, the cells were placed in a 42 °C water-bath for 90 seconds, and then immediately supplemented with 500 µl room-temperature LB medium and incubated at 37 °C for 1 hour. The cells were then isolated by centrifugation for 4 minutes at 13000 RPM. 400 µl supernatant was then removed, and the

---

[1] https://www.genomics.agilent.com/primerDesignProgram.jsp

cells were resuspended in the remaining LB medium. 100 µl of the transformed *E. coli* cells were then plated out onto LB agar ampicillin or kanamycin plates, and incubated overnight at 37 °C.

##### 1.2.4 Small Scale Plasmid DNA Purification

XL10 colonies were picked under sterile conditions from LB-agar plates, and added to 5 ml 2TY medium, supplemented with ampicillin or kanamycin, to ensure selective growth of the picked colonies. This pre-culture was then shaken at 160 rpm for 16 hours at 37 °C. The plasmid was then isolated via alkaline lysis using the QIAprep® Spin Miniprep Kit High-Yield Protocol.

##### 1.2.5 Affimer Expression

*E. coli* BL21 (DE3) cells, harbouring the pET11-Affimer-HIS6 plasmid, were collected under sterile conditions from a glycerol stock or from a LB-agar plate, and added to 5 ml LB medium together with ampicillin or kanamycin, to ensure selective growth. This pre-culture was shaken at 200 rpm for 18 hours at 37 °C, before being used to inoculate 400-1000 mL LB-media containing the appropriate antibiotic. Following inoculation, cultures were induced with 0.5 mM IPTG at OD 0.6, and incubated at 30 °C for at least six hours.

##### *1.2.6* CTB Expression from *Vibrio sp. 60*

For the production of wild-type CTB, a preculture of the *Vibrio sp.60* stock was initially set up in 100 mL high salt LB growth media, with 100 µg $mL^{-1}$ of ampicillin. This was incubated at 30 °C for 18 hours, before being used to inoculate 5 x 1 L high salt LB growth media. Once the $OD_{600}$ reached 0.6-0.8, 240 mg/L IPTG was added to the flasks to induce protein expression. Following IPTG induction, the cells were incubated at 30 °C for another 24 hours, before cell pellet isolation by centrifugation at 17000 x*g* for 25 minutes. The supernatant was retained, as this contained the protein of interest, and was thus separated from the cell pellet, and purified by ammonium sulfate precipitation.

Solid ammonium sulfate was added and dissolved to a final concentration of 57% w/v. The saturated solution was then stirred for two hours using a magnetic stirrer at room temperature. The solution was then centrifuged at 18,000 x*g* for 30 minutes, and the supernatant was discarded, whilst the pellet was resuspended in 20 mL PBS. This in turn was centrifuged at 18,000 x*g* for 10 minutes to remove insoluble material, and to allow the suspension to be filtered through a 0.8 µm Sartorius Minisart filter. Finally, the filtered suspension was separated on a lactose affinity column at 4°C, and eluted with lactose elution buffer. The eluate was dialysed 4 times against high salt PBS.

##### *1.2.7* Expression and Purification of CTB variants from *E. coli*

1 L LB media was inoculated with an overnight culture of BL21 cells transformed with the relevant pSAB-CTB plasmid and protein expression induced with 0.5 mM IPTG. The induced culture was incubated for ~20 hours, and then centrifuged at 10,000 x*g* for 10 min at 4 °C to pellet the cells. The supernatant was retained and combined with ammonium sulfate to precipitate the protein as outlined above. The presence of the surface exposed H13 in CTB confers affinity for Ni-NTA resin, offering an alternate purification route. This purification strategy was often employed when expressing multiple CTB variants, as purification by Ni-NTA chromatography could be set up in a higher throughput manner than lactose affinity chromatography. A less pure eluate was obtained via this route, necessitating purification by gel filtration.

##### 1.2.8 Cell Disruption for Protein Purification

Cells were pelleted by centrifugation for 10 minutes at 10,000 x*g.* The supernatant was discarded, and the remaining cell pellet was resuspended in a lytic cocktail containing 8 mL lysis buffer, 0.8 mL BugBuster (Novagen). This suspension was incubated at room temperature on a rocker for 40 minutes for standard proteins, or 20 minutes, followed by an additional 20 minute incubation at 50 °C, to be carried out when purifying Affimers. The lysate was then clarified by centrifugation at 10,000 x*g* for 30 minutes, followed by filtration through a 0.45 µm Sartorius filter.

##### 1.2.9 Purification of His-tagged Proteins

2 mL Ni-NTA slurry was transferred to a Bio-Rad Econo-Pac® Chromatography Column, and washed with 25 mL lysis buffer to dilute and remove the resin storage solution. The liquid was drained, and the clarified cell lysate was transferred to the column containing the resin, which was capped at both ends, and placed on a rocker for 30 minutes to facilitate binding of the His-tagged protein to the Ni-NTA resin. The lysate was drained from the column, and the settled resin was washed with 50 mL Ni-NTA Wash Buffer. The bound material was then eluted with 10 mL Ni-NTA Elution Buffer and collected in 2 mL fractions.

##### 1.2.10 Purification by Gel Filtration Chromatography

Following elution from affinity columns, protein solutions were further purified by gel filtration chromatography. All columns were stored in 20% ethanol, and all solutions to be applied directly to size exclusion columns were filtered through a 0.2 µm Sartorius Minisart filter. Columns were equilibrated initially with water followed by 1.5 column volumes of an appropriate protein buffer. When displacing the ethanol, flow rates were set at half the standard flow rate to prevent exceeding the pressure limit of the column.

For large-scale purification of proteins, a HiLoad™ 16/60 Superdex™ 75 prep-grade column (GE Healthcare) was attached to an Äkta Purifier FPLC system, and flowed at a rate of 1 mL $min^{-1}$. Protein samples ranging from 500 µl to 5 mL in volume, and between 10 kDa and 80 kDa in mass, were loaded onto this prep-grade column.

For analytical scale purification of proteins, a Superdex™ 75 Increase 10/300 column was attached to the FPLC system, and flowed at a rate of 0.5 mL $min^{-1}$. Volumes ranging from 50 µl to 1 mL were loaded onto the analytical column, with a mass range of 10 – 80 kDa.

Proteins samples for purification or analysis were injected manually into the injection loop of the FPLC system. The loop was then flushed with buffer (1.5 x the loop volume) in an automated process to apply the loop contents to the resin. Fractions were collected once the void volume of the column had been reached, and were analysed by a detector measuring absorbance at 280 nm, or 488/555 nm if purifying fluorescently-labelled proteins.

##### 1.2.11 Concentration of Protein Samples

Following size exclusion chromatography, protein samples were concentrated using Amicon Ultra-15 Centrifugal Filter Units with cut-off range of 3 - 30 kDa. 5 mL fractions were transferred to the filter units, and centrifuged at 3,500 x*g* until the protein was concentrated to an appropriate volume. As the protein collects on the filter, resuspension of the protein every 15 minutes using a pipette was necessary.

##### 1.2.12 Protein labelling

###### *1.2.12.1 Labelling Affimers with Alexa Fluor 555 C2 maleimide*

Affimers carrying a C-terminal cysteine residue were concentrated to 150 µM in TCEP phosphate buffer (50 mM phosphate, 150 mM NaCl, 0.5 mM TCEP, pH 7.2). 400 µl Affimer was combined with 0.94 mg Alexa Fluor 555 C2 maleimide (aliquoted and lyophilised) (equal to five molar equivalents). Samples were incubated at room temperature for four hours in foil, before purification using a PD-10 desalting column, concentration to 250 µl, and a final purification by gel filtration using an analytical Superdex 75 increase column. Concentration of labelled Affimer was calculated using an equation provided by Thermo Fisher Scientific:

$$[ACTA]M = \frac{(A_{280} - (A_{555} * 0.08)) * dilution\ factor}{\mathcal{E}\ cm^{-1}\ M^{-1}}$$

###### *1.2.12.2 Labelling CTB with aminooxy Alexa Fluor 488*

CTB was labelled with aminooxy Alexa Fluor 488 by oxidising the N-terminal threonine, and combining the subsequent oxidised CTB with two molar equivalents of the lyophilised fluorophore. CTB (500 µM [protamer concentration], in phosphate buffer only), L-methionine (10 equivalents), and $NaIO_4$ (5 equivalents) in phosphate buffer, were incubated together at room temperature out of light for 45 minutes, before buffer exchanging into phosphate buffer (**pH 6.8**) using a PD10 desalting column. The 1 mL eluate was then concentrated to 0.5 mL (~500 µM), and added to 0.4 mg lyophilised fluorophore (2 molar equivalents) and 4.5 µl aniline (neat). To reduce the number of fluorophores per pentamer, the amount of fluorophore can be reduced to 0.4 molar equivalents. After a 20 hour incubation, the protein was purified using a G25 desalting column, concentrated to 0.5 mL, and purified by gel filtration.

#### 1.3 Phage Display

##### 1.3.1 Preparation of ligand-coated plates

###### *1.3.1.1 Streptavidin plate preparation*

Streptavidin-coated plates were prepared by coating Nunc MaxiSorp flat-bottom 96 well plates with 100 µl 100 nM streptavidin diluted in *PBS*. The plate was sealed, and incubated at 37°C overnight to adsorb the protein to the plate surface. The following day, the 96-well plate was washed with *PBS-T*, and then blocked with *blocking buffer* overnight at 37°C. The commercial blocking buffer is casein-based, and blocks sites in the wells not occupied by streptavidin to prevent non-specific binding. After the incubation, the plate was washed once more with *PBS-T* prior to use.

###### *1.3.1.2 GM1 plate preparation*

GM1-coated plates were prepared by coating Nunc MaxiSorp flat-bottom 96 well plates with 100 µl 1.3 µM solution of GM1 ganglioside in methanol. The methanol was allowed to evaporate, thereby adsorbing the ganglioside to the plates. The GM1 plates were blocked overnight by incubating them at 37°C with a casein-based blocking buffer. The following day, the plate was washed once with 300 µl PBST on a plate washer immediately before use.

##### 1.3.2 Panning the phage library using streptavidin-coated strips

In the first step, streptavidin-coated plastic wells were prepared for pre-panning the phage-Affimer library. 300 µl *blocking buffer* was added to four wells of a Pierce streptavidin-coated 8-well strip (Thermo Scientific). The strip was sealed, and incubated overnight at 37°C. Three of the wells were to be used for pre-panning the library, with the final well required for panning the phage against the biotinylated target protein. A colony of ER2738 cells was transferred into 5 mL 2TY supplemented with 12 µg $mL^{-1}$ tetraycline (2TY tet) for overnight growth.

The wells were washed three times with *PBS-T* using a plate washer. 100 µl *blocking buffer* and 20 µl of CTB-biotin (1.5 µM/100 µg $mL^{-1}$) were added to the fourth well, for capture of the target to the streptavidin, whilst 100 µl *blocking buffer* only was added to the first three pre-panning wells. The strips were sealed once more, and incubated for two hours at room temperature on a vibrating platform shaker. Buffer from the first pre-pan well was removed, and was replaced with 100 µl *blocking buffer* and 5 µl of the phage library. The phage was then mixed and incubated in the well on a vibrating platform for 40 minutes. The buffer from the second pre-pan well was removed, and the solution from the first pre-pan well was transferred. The transferred phage was incubated for 40 minutes once more, and the same process was repeated for the third pre-pan well. This process was employed to remove phage-Affimer particles that bind to a component of the well, such as the bound streptavidin, or to other adsorbed proteins such as casein. The wells containing the target protein were washed six times, before adding the phage from the pre-pan wells. A two-hour incubation at room temperature followed on the vibrating platform shaker to allow the transferred phage to bind to the immobilised target.

##### 1.3.3 Phage elution and propagation

A fresh culture of ER2738 *E. coli* cells was grown to an $OD_{600}$ of approximately 0.6 by diluting the overnight culture 1/15 into 8 mL 2TY, and incubating the solution at 37°C for one hour.

Whilst the culture was growing, the panning well was washed 27 times with *PBS-T*. The phage was then eluted by adding 100 µl of 0.2 M glycine, pH 2.2, and incubating the solution for ten minutes at room temperature before neutralisation with 15 µl of 1M Tris-HCl, pH 9.1. The resulting solution was transferred immediately to the fresh 8 ml culture of the ER2738 *E. coli* cells. To ensure that all bound material had been successfully eluted, 100 µl of triethylamine (1.4% in *PBS*) was added to the well, and left for six minutes. 50 µl 1M Tris-HCl, pH 7.0 was added to neutralise the triethylamine, and the solution was transferred to the 8 mL culture. The culture was placed in a 37 °C incubator for one hour, and shaken once, by hand, after 30 minutes.

1 µl of the culture was combined with 100 µl 2TY, and plated out onto a LB-agar carbenicillin plate. The remainder of the culture was centrifuged to pellet the cells, and resuspended into 100 µl media. The entire suspension was transferred and spread across a second LB-agar ampicillin plate. The plates were incubated at 37 °C overnight. The following day, the colonies on the plate containing 1 µl of cells were counted, and multiplied by 8,000 to determine the total number per 8 ml of cells. Successful panning rounds typically generated between 0.5 – 2 x $10^6$ cells.

5 mL 2TY supplemented with 100 µg/ml carbenicillin (2TY carb) was added to the plate containing the full culture suspension, and a plastic spreader was used to scrape the cells into the solution. The 5 mL was transferred to a falcon tube, and a further 2 mL 2TY was used to scrape any remaining cells from the plate surface. The $OD_{600}$ was measured of a 1:10 dilution, and the suspension was diluted into 2TY to make an 8 mL culture of $OD_{600}$ = 0.2. The dilution was incubated at 37 °C, 230 rpm, for one hour.

0.32 µl M13K07 helper phage (titre ca. $10^{14}$/ml) was added to the culture after one hour, and left to incubate at 37 °C, 90 rpm, for 30 mins. The helper phage also infect the bacteria, providing the phage-Affimer with additional components to form infectious particles. The helper phage also confers resistance to kanamycin. To select only the bacteria harbouring phage-Affimer particles and the helper phage, 16 µl of kanamycin (25 mg $ml^{-1}$) was added, and the culture was incubated overnight at 25 °C, 170 rpm.

The phage-infected cultures were centrifuged at 3,500 *xg* for 10 mins to pellet the infected cells, leaving the newly formed phage-Affimer particles in solution. The supernatant was retained and transferred to fresh tubes. 125 µl phage-containing supernatant was retained for the next panning round.

##### 1.3.4 Phage purification and storage

To purify the phage, 2 ml of PEG-NaCl precipitation solution (20% (w/v) PEG 8000, 2.5M NaCl) was added to the phage-containing supernatant, mixed, and incubated overnight at 4 °C. The following day, the precipitated phage were pelleted at 4,816 *xg* for 30 mins, and the supernatant was discarded. The pellet was resuspended in 320 µl TE buffer, transferred to a 1.5 mL Eppendorf tube, and centrifuged at 16,000 *xg* for 10 mins. The supernatant contained purified phage, which could be used directly in a subsequent panning round, or could be combined with 50% glycerol and stored at -80 °C for long-term storage.

##### 1.3.5 Panning the phage library using streptavidin-coated magnetic beads

After propagation of the phage from the first panning round, a second panning round was carried out to expose the phage to the same target, but presented in a different manner. Rather than exposing the phage to CTB-biotin captured onto streptavidin-coated plastic wells, the biotinylated target would be presented on streptavidin-coated magnetic beads. This difference in presentation should minimise the number of background binders carried through the screen by presenting a very different binding surface.

As in the first round, a colony of ER2738 *E. coli* cells was cultured overnight in 5 mL 2TY tet. 20 µl streptavidin beads (Dynabeads MyOne Streptavidin T1) were combined with 100 µl *blocking buffer* and incubated overnight at room temperature on a Stuart rotator.

The next day, the beads were centrifuged at 800 *xg* for one minute, placed onto a magnetic Eppendorf rack, and resuspended in 100 µl fresh *blocking buffer*. The magnetic rack draws the beads to the side of the tube, allowing buffer to be changed without losing the streptavidin beads. 125 µl of the phage retained from the initial propagation step, or 5 µl of the purified phage, was combined with blocking buffer to a total volume of 250 µl in Eppendorf LoBind Tubes. 25 µl of the pre-blocked streptavidin beads was added to the phage, and incubated on a Stuart rotator for one hour. Beads were separated from the phage by centrifugation and placement on the magnetic rack. The supernatant was transferred to a tube containing another 25 µl of pre-blocked streptavidin beads, and the pre-panning process was repeated.

15 µl CTB-biotin (1.5 µM) was combined with 200 µl blocking buffer, and 50 µl streptavidin beads, before being placed on a Stuart rotator for 30 minutes to bind. The beads were washed three times with 500 µl blocking buffer by removing the buffer, resuspending the beads, and then immobilising the beads to the magnetic rack, removing the buffer, and iterating the process. The supernatant containing the pre-panned phage was then transferred to the beads containing the captured CTB-biotin, and placed on a Stuart rotator for 45 minutes. After the incubation, the beads were washed manually 27 times with PBS-T.

A fresh 8 mL culture of ER2738 *E. coli* cells was set up, and the bound phage were eluted and used to infect the culture as described above. Plates were prepared and the phage propagated and stored as described previously. 200 µl phage supernatant was retained for the following panning round.

##### 1.3.6 Panning the phage library against GM1-bound CTB

CTB is a lectin that binds with high affinity and specificity to GM1 ganglioside. The phage display protocol could therefore be modified to screen the phage library against GM1-bound unmodified CTB. GM1 plates were prepared, blocked, and washed as described in *GM1 plate preparation.* An overnight culture of ER2738 *E. coli* cells was prepared.

10 µl purified phage from the second panning round (CTB-biotin bound to streptavidin-coated magnetic beads) was added to 190 µl blocking buffer, and was pre-panned against four GM1-coated wells. 50 µl 1 µM solution of CTB was captured onto the GM1-coated surface of the fifth well, whilst the sixth well was retained as a negative control. Meanwhile, two 5 mL ER2738 *E. coli* cell cultures were prepared by diluting the overnight culture 1:15, and incubating the solution at 37 °C for one hour, 230 rpm. The panning wells were washed six times with PBS-T, before addition of the phage from the third pre-panning well, which was split between the two panning wells. After a one hour incubation, phage were eluted, and used to infect the two 5 mL ER2738 *E. coli* cell cultures. The Plates containing single colonies were taken forward for analysis by phage ELISA.

##### 1.3.7 Phage ELISA

An enzyme-linked immunosorbent assay (ELISA) was carried out to validate the binding of a phage-Affimer subset to a screened target.

200 µl aliquots of 2TY carb were transferred to a 96-well V-bottom deep well plate using a multichannel pipette. From the final panning round, 24 - 48 individual colonies were picked and used to inoculate the wells. The plate was then incubated overnight at 37°C, 1050 rpm. 200 µl 2TY was then transferred to the wells of a fresh deep well plate, which were inoculated with 25 µl of the overnight culture, before being incubated in turn for one hour at 37°C, 1050 rpm.

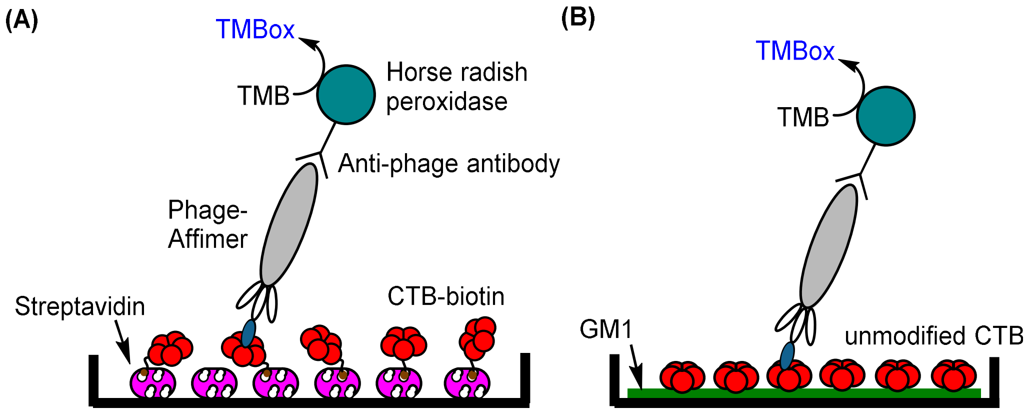

*Supplementary Figure 7. Phage ELISA schematic. (A) CTB-biotin is immobilised to streptavidin-coated plates, and presented to a phage-Affimer species. The bound phage is detected by anti-phage antibody, conjugated to horse radish peroxidase (HRP). Binding is detected by the addition of TMB and hydrogen peroxide, which is converted to a blue product by the peroxidase. The blue product absorbs at 620 nm and can be detected by plate reader. (B) Unmodified CTB is immobilised to GM1 ganglioside-coated plates, and presented to a phage-Affimer species. Detection of bound phage is facilitated through use of a peroxidase-conjugated antibody, and the peroxidase substrates TMB and hydrogen peroxide.*

After one hour, 5 µl M13K07 helper phage (titre ca. $10^{14}$/ml) was diluted into 5 ml 2TY carb, and 10 µl of the diluted stock was transferred into each of the wells of the one-hour culture. This culture was incubated for 30 minutes at room temperature, 450 rpm. The rpm was reduced to enable efficient infection of the cells with the helper phage, which was required in turn to supplement the phage-Affimer present in the infected cells with various complement proteins required for infectious phage-Affimer particles to escape into the surrounding media.

Finally, kanamycin was added to the cultures by diluting the stock 20 fold into 2TY, and transferring 10 µl of the diluted antibiotic into each of the wells. As the helper phage confers kanamycin resistance to infected bacteria, this would effectively kill any *E. coli* cells that had not taken up the helper phage. This was incubated overnight at room temperature, 750 rpm. The phage-infected culture was centrifuged at 3500 *xg* for ten minutes to pellet the bacteria, leaving the Affimer-expressing phage in the supernatant. The supernatant could then be used directly in the ELISA.

Plates were prepared with either streptavidin or GM1 ganglioside. To half of the plate, CTB-biotin (1 µM), or unmodified CTB (1 µM), was added and allowed to bind for one hour, before being washed three times with PBS-T to remove unbound material.

40 µl of phage-containing supernatant was then transferred to the first and seventh well of the plate such that each phage was tested against the target, as well as the negative control well. 10 µl undiluted 10x Casein Blocking Buffer was added to supplement the phage. Following a one hour incubation, the plate was washed 27 times with PBS-T.

Commercially acquired Anti-Fd-Bacteriophage HRP was diluted 1000 fold in blocking buffer, and 50 µl was added to each well. After another one hour incubation, the plate was washed 12 times with PBS-T. Binding was assayed by adding 50 µl SeramunBlau solution, which contained the TMB and hydrogen peroxide. The plate was allowed to develop for approximately four minutes, before the absorbance at 620 nm was measured using a plate reader.

#### 1.4 Biophysical Techniques

##### 1.4.1 Isothermal Titration Calorimetry

Isothermal titration calorimetry (ITC) experiments were carried out to collect information on the thermodynamics of CTB-Affimer interactions.

The $iTC_{200}$ Microcalorimeter User's Manual[2] is a useful tool for practical understanding of how to carry out ITC experiments, and provides detailed information on instrument navigation.

Extensive dialysis of both proteins was required to match buffers before running experiments. All ITC experiments were performed using a MicroCal $ITC_{200}$. Water in the reference cell was replaced at the beginning of a set of experiments, using a 1 mL Hamilton syringe. The sample cell was then washed extensively with PBS-T to ensure that no protein remained from any previous experiments. The cell was additionally washed with water, followed by the experiment buffer, to remove any bubbles remaining from the detergent used in the previous wash step, and to equilibrate the sample cell. 300 µl protein 'receptor' was gently drawn up into a 1 mL Hamilton syringe, and purged from any bubbles. The sample cell was then gently filled. The tip of the syringe was then drawn up to the ledge formed at the interface between the sample cell, and the overflow reservoir. Any excess liquid was removed up to this point, thereby ensuring that the volume was standardised across all experiments.

The titration syringe was then loaded with ligand from a microcentrifuge tube, and the fill port was closed. The titration syringe was then placed back into the tube, and the titration syringe was purged, refilled, and then transferred into the sample cell. The calorimeter was heated to 25 °C or 35 °C before carrying out titrations.

An initial sacrificial titration of 0.5 µl was required, before 19 x 2 µl titrations were carried out, each spaced out over 4 seconds, with a 120 second delay in between each injection. Experiments were initially carried out at 100 µM CTB, titrated into 10 µM Affimer. These concentrations were then varied, if necessary, to obtain a sigmoidal shape of curve. For each set of concentrations, a CTB titration into buffer was carried out to obtain a measure of the heat of dilution of CTB into buffer. These data could then be subtracted from the data obtained from ligand into receptor titrations to bring the baseline closer to zero, thereby facilitating curve fitting. Data were analysed and fitted with a 'one site bind' model using Origin Microcal software.

---

[2] http://www.biophysics.bioc.cam.ac.uk/files/iTC200_User_Manual_Rev_D.pdf

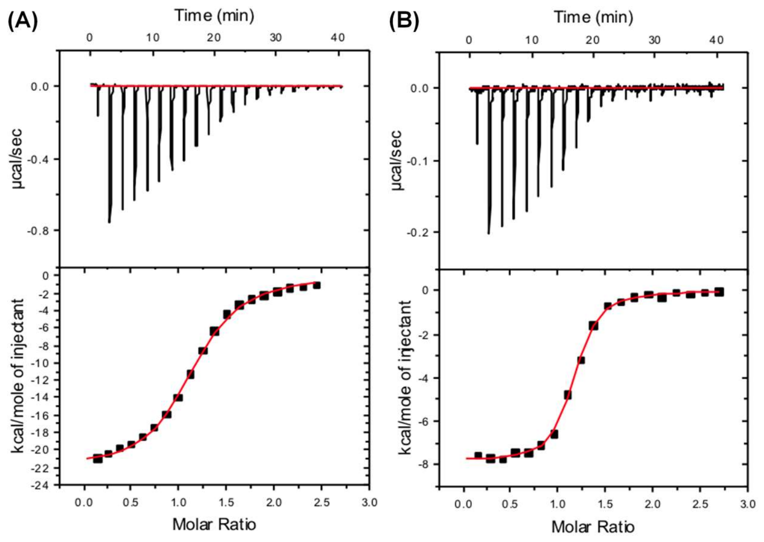

Supplementary Figure 8. Results of ITC experiments (A) Thermogram and binding isotherm generated following the titration of 100 µM CTB into 10 µM C6 at 35$^{o}$C by ITC. The data are fitted with a one-site binding model using Microcal Origin software following baseline subtraction using CTB into buffer titrations. (B) Thermogram and resultant Wiseman binding isotherm following high concentration titrations of 200 µM CTB into 17 µM A2 at 35$^{o}$C by ITC. The data are fitted with a one-site binding model using Microcal Origin software following baseline subtraction using CTB into buffer titrations.

##### 1.4.2 Affimer-lectin binding assay

An Affimer-lectin binding assay (ALBA) was developed to validate the binding of Affimers to CTB (**Error! Reference source not found.**).

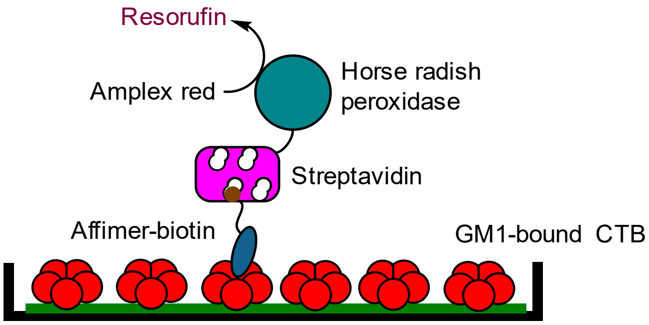

*Supplementary Figure 9. Schematic for an Affimer-lectin binding assay. A GM1-binding lectin, in this case CTB, is captured onto a GM1-coated surface. A biotinylated Affimer binding CTB is recognised by a streptavidin-peroxidase conjugate, which is quantified by addition of Amplex Red and hydrogen peroxide to form the fluorescent resorufin.*

In order to carry out this assay, biotinylated Affimers were required. Affimers carrying a C-terminal cysteine were expressed, and biotinylated using biotin-PEG2-maleimide. Affimers were concentrated to 100 µM, and combined with a 20-fold molar excess of biotin-PEG2-maleimide in *phosphate buffer* supplemented with 1 mM TCEP. After an overnight incubation at room temperature, the Affimer was purified by using a PD-10 desalting column, or by dialysis into *phosphate buffer*.

50 µl GM1 ganglioside in methanol (1.3 µM) was added to wells (as required) of a Greiner Bio-one high binding black 96 well plate. The plate was then left in a lamina-flow hood until the methanol had completely evaporated. To block the plate and occupy any remaining adsorption sites, *blocking buffer* was added at a volume of 300 µl per well. The plate was covered, and incubated overnight at 37°C. Following incubation,

the blocking solution was removed, and the plate was washed three times with *PBS-T*. A Combi Reagent Dispenser and standard tube dispensing cassette were used to facilitate accurate and rapid liquid transfer into the wells of the plate.

After the wash steps, 50 µl lectin (1 µM) was transferred to the appropriate wells, and left to bind for one hour at room temperature. Half the wells were deliberately left free of CTB to act as a control. The wash step was then repeated to remove any unbound CTB, and subsequently 100 µl biotinylated Affimer (250 nM) was added to each well. The Affimer was in turn allowed to bind for one hour before repeating the wash step a third time. In order to detect the presence of bound Affimer, 50 µl of a streptavidin-HRP complex (Ultra Streptavidin-HRP, Life Technologies), diluted at 1:1000 in *phosphate buffer*, was added to the wells of the plate. The one hour incubation and wash step was repeated, with an additional four washes, before one final rinse with *PBS* or *phosphate buffer*. The substrate used for peroxidase detection was a solution of Amplex Red and hydrogen peroxide in *PBS*, both at a final concentration of 5 µM. 50 µl of this substrate was added per well, and the plate was loaded into the plate reader for fluorescence detection at 585 nm.

#### 1.5 Tissue Preparation and Analysis

##### 1.5.1 Mammalian Cell Culture

###### *1.5.1.1 Passage of cells*

Vero cells were cultured in *cell culture media*, a supplemented Dulbecco's Modified Eagle's Medium (DMEM) containing foetal bovine serum, added due to the presence of essential growth factors for the cell proliferation, and a combination of antibiotics penicillin and streptomycin, to prevent the bacterial and fungal infections. Cultures were grown in T75 cell culture flasks at 37 °C with 5% $CO_2$ until 90% confluency, approximately after three days of culturing. At this stage, cells were washed once with 10 mL sterile *PBS*, and combined with 3 mL TrypLE Select (Life Technologies), a trypsin-based protease solution to detach cells from the flask surface. After 5 minutes at 37 °C, flasks were placed under a light microscope to verify complete detachment. If large numbers were seen to still be adhering to the surface, the flask was firmly tapped, and placed in the 37°C incubator for a further 3 minutes. 7 mL Cell Culture Media was added, neutralizing the trypsin, and the detached cells were transferred to a 15 mL falcon tube. The flask was washed twice with 10 mL PBS, and 20 mL Cell Culture Media was added, together with 1 mL of the resuspended cells. Cells were then returned to the incubator for culturing at 37 °C with 5% $CO_2$.

###### *1.5.1.2 Cell seeding*

For cell-based experiments, cells were 'seeded' onto 18 mm glass coverslips in 12-well plates. After passaging cells, the falcon tube containing the cells was retained, and used for cell seeding. A Neubauer Improved haemocytometer was used for cell counting by making a 1:10 dilution of the cells, and transferring 5 µl of the diluted sample onto the haemocytometer. A coverslip was then used to draw the cells onto the central grid by capillary action. A 5x5 grid at the centre of the haemocytometer was used for cell counting, and cells from five of the boxes were counted. This number was divided by two, and multiplied by $10^6$ to calculate the number of cells per mL. For a 12-well plate, $1.2 \times 10^5$ cells per well were plated out. Cells were therefore diluted to $1.2 \times 10^5$ cells $mL^{-1}$, and 1 mL was transferred into the wells of a sterile 12-well plate, containing an 18 mm coverslip in each well. The plate was then incubated overnight at 37 °C with 5% $CO_2$ for use the following day.

###### *1.5.1.3 Cell treatment with protein*

Seeded cells in 12-well plates to be treated with protein were washed, by replacing the Cell Culture Media with 1 mL fresh media. Protein solution was then added directly to the media, and the cells were incubated for 4-6 hours at 37 °C, 5% $CO_2$ prior to cell fixation.

###### *1.5.1.4 Cell fixation*

Following treatment with protein, cells were exposed to a strong acid, to remove any surface-bound CTB, and then 'fixed', by adding a harsh cross-linking agent, effectively killing the cells and freezing them in a particular state.

Growth medium was removed, and the cells were washed three times with PBS. 0.5 mL Cell Cleansing Buffer was added, and incubated at 4 °C for 6 minutes. The cleansed cells were then washed an additional three times, before addition of 4% v/v paraformaldehyde (PFA) in PBS for 30 minutes at room temperature in a fume hood. The PFA was removed, and the cells were washed three times with PBS. Cells were then permeabilised with 0.1% Triton X-100 in PBS for one hour at room temperature in preparation for immunohistochemistry.

##### 1.5.2 *In Vivo* Experimentation

All handling of live animals was carried out either by Dr Jessica Haigh, or Professor Jim Deuchars.

###### *1.5.2.1 Animals*

All animal experimentation was carried out under a Home Office License, and in accordance with the regulations of the UK Animals, Scientific Procedures, Act 1986. Experiments were performed on young male and female C75BL/6 adult mice, bred in house.

###### *1.5.2.2 Intramuscular administration of protein*

The tongue was targeted for intramuscular injections of protein and protein complexes, since it provided a route to deliver material into the motor neurones of the brainstem. Protein was prepared at a concentration of 50 μM in phosphate buffer, before lyophilisation for reconstitution prior to injection at a concentration of 200 μM.

Paralingual injections were carried out by heavily sedating mice with isoflurane, protruding the tongue from the mouth using forceps, and injecting 2 μl of protein material using a glass micropipette mounted onto a 10 μl Hamilton syringe. Mice were perfused after 24 – 72 hours.

###### *1.5.2.3 Transcardial perfusion for tissue fixation*

Mice were deeply anaesthetised by injection of sodium pentobarbitone (60 mg $kg^{-1}$) into the abdominal cavity. To ensure that the mice were fully sedated, pedal reflexes were tested. The abdomen of the animal was then dissected transversely, and a thoracotomy was carried out to reach the heart. The left ventricle of the heart was pierced, and a blunt needle was inserted and retained in position with a clip. The right atrium was then cut, and a phosphate solution (0.1 M $NaH_2PO_4$, pH 7.4) was applied through the needle to flush the circulatory system. Perfusion was then carried out by replacing the phosphate solution with approximately 150 mL 4% PFA.

###### *1.5.2.4 Dissection and Tissue Preparation*

The brainstem was removed by dissection of the spinal cord, opening of the skull, and separation of the meninges from the brain. The brain was then removed from the skull, ensuring not to disturb the brainstem and upper half of the spinal cord. The brain and spinal cord was then transferred into a 50 mL falcon tube with 4% PFA, and kept for 24 hours at 4°C. The PFA was then removed and replaced with PBS, and the brainstem was separated from the hind brain and spinal cord for sectioning. The brainstem could then be sectioned. All sections were taken transversely on a VT1000S vibrating microtome at 50 μm thickness. The sectioned slices were collected and transferred into PBS in a 24-well plate.

##### 1.5.3 Immunohistochemistry

Tissue sections, or cultured cells, were permeabilised in 1 mL *solubilisation buffer* for one hour at room temperature. The primary antibody (e.g. rabbit anti-CTB, chicken anti-GFP) was diluted 1:1000 in *solubilisation buffer*, and 0.5 mL was applied to the tissue/cells. After an overnight incubation at 4 °C, the wells were washed three times with PBS, and a 1:1000 dilution of a fluorescent secondary antibody (Alexa Flour 488 or 555) in PBS was made and 0.5 mL added to the sample. A one hour incubation at room temperature ensued, followed by three more washes with PBS. The sample was then air-dried onto a glass microscope slide, before addition of 10 µl of Vectashield Mounting Medium, and protection of the sample with a cover slip ready for visualisation. To stain the nuclei in a sample, Vectashield Mounting Medium with DAPI was used. This medium contains DAPI stain, a fluorescent stain that tightly associates to AT-rich regions of DNA in the nuclei of cells, thus facilitating visualisation of the nuclei. Microscope slides were covered with foil, and kept overnight at room temperature to dry, before being sealed with a generic nail varnish.

##### 1.5.4 Confocal Microscopy

For confocal analysis of samples, an Axio Imager Z2 LSM880 upright confocal microscope (Zeiss) equipped with a 405 nm diode laser, Argon 458/488/514 nm, DPSS 561 nm and HeNe 633 nm lasers and a GaAsP detector. Images were captured using the Zeiss LSM Image browser software, and processed using Zen lite 2.3 software.

For wide-field analysis of samples, a Nikon Eclipse E600 microscope equipped with epifluoresence and Q-Imaging Micropublishing 5.0 camera was used. Images were captured using AcQuis image capture software.

#### 1.6 Plasmid Sequences

El Tor CTB was expressed from *Vibrio sp.60* clones harbouring the pATA13 plasmid.[1]

Affimer-pIII fusion proteins were expressed from ER2738 *E. coli* cells harbouring the pBSTG plasmid.[2]

CTB variants were expressed from a plasmid based on pMal-p5x entitled pSAB.[3]

##### 1.6.1 CTB/LTB protein sequences

Protein sequences for LTBh, LTBh T80A, El Tor CTB, and CTB A80T. Highlighted in yellow is residue A80.

| | | | | |
|---|---|---|---|---|
| **LTBh (H74-114)** | APQSITELCS | EYHNTQIYTI | NDKILSYTES | MAGKREMVII |
| **LTBh T80A** | APQSITELCS | EYHNTQIYTI | NDKILSYTES | MAGKREMVII |
| **CTB (El Tor)** | TPQNITDLCA | EYHNTQIYTL | NDKIFSYTES | LAGKREMAII |
| **CTB A80T** | TPQNITDLCA | EYHNTQIYTL | NDKIFSYTES | LAGKREMAII |
| **LTBh (H74-114)** | TFKSGATFQV | EVPGSQHIDS | QKKAIERMKD | TLRITYLTET |
| **LTBh T80A** | TFKSGATFQV | EVPGSQHIDS | QKKAIERMKD | TLRITYLTEA |
| **CTB (El Tor)** | TFKNGAIFQV | EVPGSQHIDS | QKKAIERMKD | TLRIAYLTEA |
| **CTB A80T** | TFKNGAIFQV | EVPGSQHIDS | QKKAIERMKD | TLRIAYLTET |
| **LTBh (H74-114)** | KIDKLCVWNN | KTPNSIAAIS | MEN | |
| **LTBh T80A** | KIDKLCVWNN | KTPNSIAAIS | MEN | |
| **CTB (El Tor)** | KVEKLCVWNN | KTPHAIAAIS | MAN | |
| **CTB A80T** | KVEKLCVWNN | KTPHAIAAIS | MAN | |

###### *1.6.1.1 ACTA protein sequences*

Binding loop sequences are shown in red.

**ACTA-A2**
GVGASAATGV RAVPGNENSL EIEELARFAV DEHNKKENAL LEFVRVVKAK EQQHERSHWV
DTMYYLTLEA KDGGKKKLYE AKVWVKHNQF FDYFINFKEL QEFKPVGDAA AAHHHHHHHH

**ACTA-A2-cys**
GVGASAATGV RAVPGNENSL EIEELARFAV DEHNKKENAL LEFVRVVKAK EQQHERSHWV
DTMYYLTLEA KDGGKKKLYE AKVWVKHNQF FDYFINFKEL QEFKPVGDAC AAAHHHHHHH
H

**ACTA-C6**
ASAATGVRAV PGNENSLEIE ELARFAVDEH NKKENALLEF VRVVKAKEQM DLNAGLPRTM YYLTLEAKDG
GKKKLYEAKV WVKQGLKKLK FTNFKELQEF KPVGDAAAAH HHHHHHH***

**ACTA-C6-cys**
ASAATGVRAV PGNENSLEIE ELARFAVDEH NKKENALLEF VRVVKAKEQM DLNAGLPRTM YYLTLEAKDG
GKKKLYEAKV WVKQGLKKLK FTNFKELQEF KPVGDACAAA HHHHHHHH**
